# CLASHub: an integrated database and analytical platform for microRNA-target interactions

**DOI:** 10.1101/2025.08.05.668543

**Authors:** Lu Li, Peike Sheng, Nicholas M. Hiers, Tianqi Li, Acadia L Grimme, Yuzhi Wang, Conner M. Traugot, Mingyi Xie

**Author notes:** Department of Molecular, Cellular and Developmental Biology, Yale University, New Haven, CT, 06511, USA. Department of Pharmacology, University of California San Diego, La Jolla, CA, 92093, USA.

## Abstract

MicroRNAs (miRNAs) are ∼22-nucleotide RNAs that regulate gene expression, critical for development and disease. Residing in Argonaute (AGO) proteins, miRNAs target messenger RNAs via complementary base-pairing. Current miRNA-target databases rely on indirect data from AGO crosslinking immunoprecipitation (AGO-CLIP). In contrast, CLASH (Crosslinking, Ligation, and Sequencing of Hybrids) employs proximity ligation within AGO complexes, providing direct miRNA-target interaction evidence. Existing CLASH datasets remain limited to few human and mouse samples. CLASHub integrates CLASH-defined interactions with gene and miRNA expression data from human, mouse, *Drosophila*, and *C. elegans*, covering over 20 cell types and tissues. The datasets also include samples with ZSWIM8-knockout, an essential component in target RNA-directed miRNA degradation (TDMD)—a specialized mechanism for miRNA turnover triggered by basepairing with specific target RNA—thereby providing unique insights into miRNA turnover mechanisms. CLASHub also features a user-friendly Analyzer interface for CLASH, RNA-seq, miRNA-seq, and cumulative fraction curve analysis. Leveraging these tools, we uncovered a novel TDMD “trigger” for miR-335-3p degradation in *ATP6V1G1*’s 3ʹ UTR. Thus, CLASHub offers an online platform for miRNA research, enabling discovery of novel miRNA targets and triggers across diverse contexts.

**Graphical abstract:** 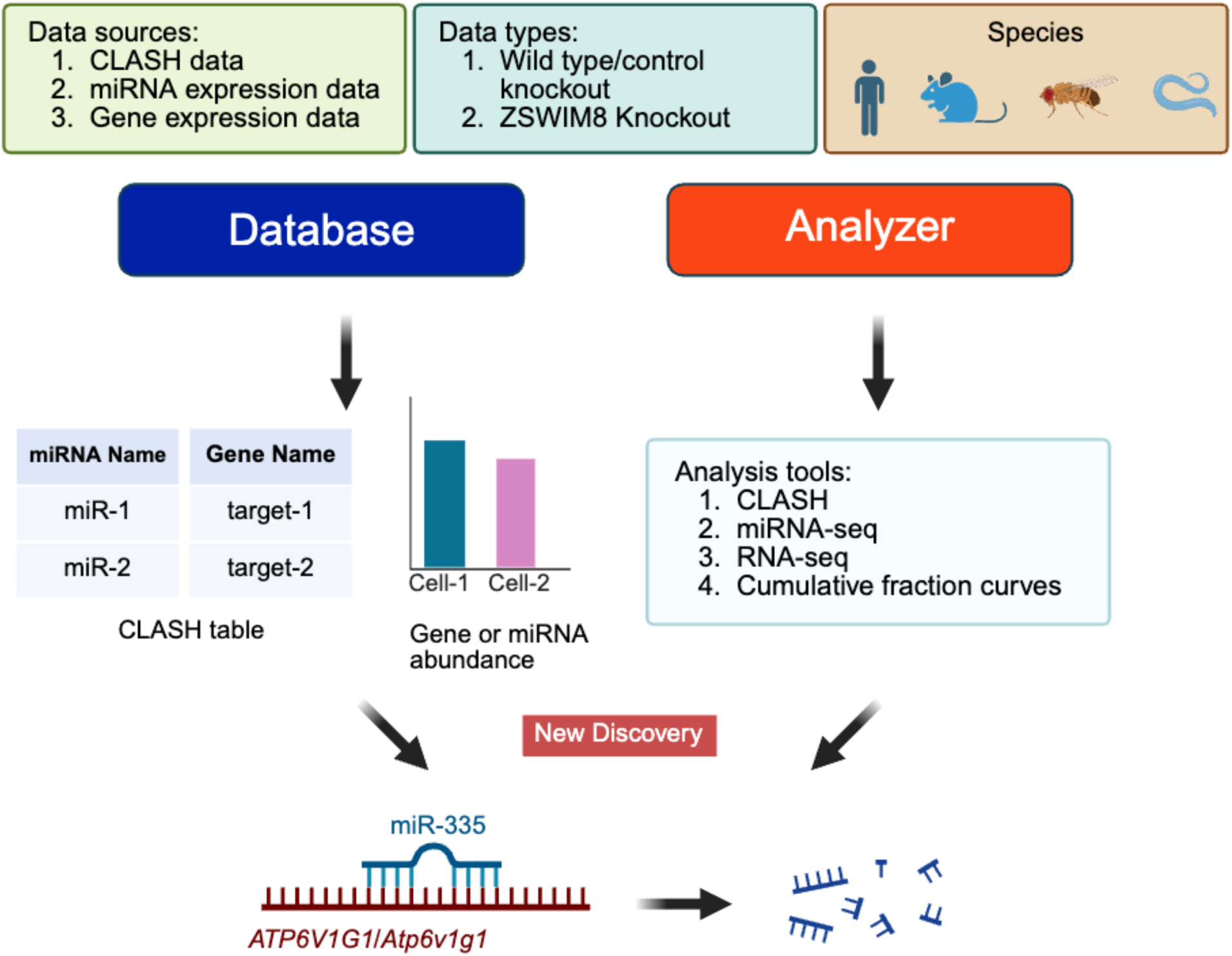

## Introduction

MicroRNAs (miRNAs) are small non-coding RNA molecules, 20–24 nucleotides in length, that play pivotal roles in post-transcriptional gene regulation in eukaryotic cells (Kim et al., 2025). They function by guiding the miRNA effector protein Argonaute (AGO) to complementary sequences within target RNAs – most commonly within the 3ʹ untranslated regions (3ʹ UTRs) of messenger RNAs (mRNAs) - to induce mRNA degradation or inhibit translation, thereby fine-tuning gene expression (Bartel, 2018). While miRNA target sites can also be found in coding sequences (CDSs) and 5′ UTRs, these are generally less frequent or less effective than those in the 3′ UTR (Hausser et al., 2013; Lytle et al., 2007). Through this regulatory mechanism, miRNAs control a multitude of cellular processes such as development, differentiation, proliferation, and apoptosis (DeVeale et al., 2021; Nassim Mahtal et al., 2022; N. Mahtal et al., 2022; Santovito & Weber, 2022). Consequently, aberrations in miRNA expression or function can disrupt these critical processes, contributing to the pathogenesis of various diseases, including cancer, neurodegenerative disorders, and cardiovascular conditions (Cui et al., 2024).

Understanding miRNA-target interactions is fundamental to unraveling the complexities of gene regulatory networks. High-throughput sequencing approaches, particularly crosslinking immunoprecipitation (CLIP)-based techniques such as HiTS-CLIP and PAR-CLIP have provided valuable insights into miRNA-binding sites genome-wide (Anastasakis et al., 2021; Chi et al., 2009; Hafner et al., 2010; O’Connor et al., 2021). These methods rely on ultraviolet (UV) light-induced crosslinking of RNA-protein complexes, immunoprecipitation of AGO-bound RNAs, followed by sequencing to identify miRNA-interacting RNA fragments. However, CLIP-based methods identify AGO binding sites on target RNAs but do not directly reveal which specific miRNAs are engaged at those sites. To overcome this limitation, the CLASH (crosslinking, ligation, and sequencing of hybrids) approach was developed, which resembles AGO-CLIP protocols but with an additional miRNA-target proximity ligation step to physically join miRNAs directly to their interacting target RNAs (Gay et al., 2018; Helwak & Tollervey, 2014). These resultant miRNA-target chimeras or “hybrids” are then sequenced to generate direct evidence of specific miRNA-target interactions and their exact binding sites. Thus, CLASH significantly enhances the accuracy and resolution of miRNA regulatory network analyses.

Despite advances in experimental techniques, challenges remain in the comprehensive identification and analysis of miRNA-target interactions across diverse species and cellular contexts. Existing databases such as miRTarBase (Huang et al., 2022), TargetScan (McGeary et al., 2019), and miRDB (Chen & Wang, 2020) compile predicted and experimentally validated miRNA targets but often lack cell type-specific information and direct evidence of interactions. Moreover, few resources integrate data from multiple organisms or provide tools for analyzing how changes in miRNA expression influence target gene expression globally (Chang et al., 2020; Hauschild et al., 2023). To address these challenges, CLASHub 1.0 (hereafter, CLASHub) was developed as a comprehensive and user-friendly platform that integrates a rich database of CLASH-derived miRNA-target interactions. CLASHub aggregates and standardizes CLASH data from diverse studies, incorporating both newly generated and previously published datasets. The database includes CLASH data from four species—*Homo sapiens* (human), *Mus musculus* (mouse), *Drosophila melanogaster* (fruit fly) and *Caenorhabditis elegans* (nematode)—spanning a variety of cell types and tissues. In addition to CLASH-derived miRNA-target interaction data, CLASHub integrates gene and miRNA expression profiles obtained from publicly available datasets, allowing for an informed understanding of miRNA-target interactions with RNA abundance.

Beyond serving as a comprehensive database, CLASHub provides powerful analytical tools that extend its functionality. For CLASH analysis, users can upload raw sequencing files to systematically identify miRNA-target RNA hybrids. The web portal would return a detailed table specifying miRNA-target pairs, genomic coordinates, base pairing patterns, target conservation scores, RNA types (e.g., mRNA, ncRNA), transcript location (e.g., 3ʹ UTR, CDS), and hybrid abundance. The integrated RNA-sequencing (-seq) and miRNA-seq analysis modules allow users to quantify global gene expression and miRNA changes, facilitating functional validation of miRNA targets identified by CLASH. Furthermore, direct evaluation of the global impact of specific miRNAs on target gene expression can be achieved by uploading differential gene expression results. These results can then be used to produce cumulative fraction curves that detail changes in miRNA target repression, utilizing target predictions from both CLASH-derived and TargetScan datasets. Collectively, these integrated analyses streamline discovery and validation, significantly enhancing the efficiency and depth of miRNA research.

Recently, there have been many advances in the subject of miRNA turnover across various animal and cellular models, primarily through a mechanism termed target-directed miRNA degradation (TDMD) (Ameres et al., 2010; Cazalla et al., 2010; Han et al., 2020; Jones et al., 2023; Li et al., 2021; Sheng et al., 2023; Shi et al., 2023; Shi et al., 2020). In TDMD, specific target RNAs contain specialized miRNA complementary binding sites - referred to as TDMD “triggers” within 3ʹ UTRs or non-coding RNAs, that induce rapid degradation of interacting miRNAs (Li et al., 2025). This process is thought to be catalyzed by trigger-mediated AGO conformation change and subsequent recruitment of the Culin-RING E3 ubiquitin ligase-ZSWIM8, Dora, or EBAX-1 in vertebrates, *Drosophila*, and *C. elegans*, respectively (Han et al., 2020; Sheu-Gruttadauria et al., 2019; Shi et al., 2020). The ZSWIM8-containing complex then catalyze AGO ubiquitination for proteasomal decay and subsequent miRNA degradation, thereby reducing miRNA abundance and activity. We have recently demonstrated the utility of CLASH data in the screening and identification of novel TDMD triggers, having validated 7 of the 11 currently accepted endogenous triggers in animals (Hiers et al., 2024; Li et al., 2021; Sheng et al., 2023). Given the platform’s comprehensive integration of diverse datasets, CLASHub is uniquely suited to investigate specific miRNA turnover processes like TDMD. Notably, the platform includes newly added datasets from ZSWIM8-knockout cells and tissues, which were not previously available (Table S1-S3) (W. L. Bullard et al., 2019; Fields et al., 2021; Gay et al., 2018; Li et al., 2025; Manakov et al., 2022; Moore et al., 2015; Sheng et al., 2023). Leveraging CLASHub, we identified a novel TDMD trigger for miR-335-3p degradation. These findings highlight its value as a resource for discovering novel miRNA regulatory interactions. The platform is publicly accessible at https://clashub.rc.ufl.edu/.

## Results

### CLASHub Databases

#### Overview of dataset composition in CLASHub

CLASHub integrates three main data types for studying miRNA regulation: CLASH data, gene expression data (RNA-seq), and miRNA expression data generated using Accurate Quantification of miRNA by Sequencing (AQ-seq), which improves the accuracy of miRNA quantification by minimizing adapter ligation bias during small RNA library preparation (H. Kim et al., 2019). These datasets were derived from publicly available sources and newly generated data in this study (Figure 1A). In contrast to previously reported CLASH data, which mainly focus on wild-type samples, our newly generated datasets significantly expand on ZSWIM8-knockout conditions. These datasets span four species—human, mouse, *Drosophila*, and *C. elegans*—and include 51 control samples (wild-type or non-targeting sgRNA) and 39 knockout samples targeting ZSWIM8 orthologs: *ZSWIM8* in humans, *Zswim8* in mice, *Dora* in *Drosophila*, and *ebax-1* in *C. elegans*. The samples originate from 17 different cell lines or tissues, 15 of which have not been previously reported (Figures S1A–B). All databases are built on an abundant number of datasets and details of cell lines, tissues, and sample sources are summarized in Figures 1B-D, S1 and Table S1-3.

**Figure 1.**
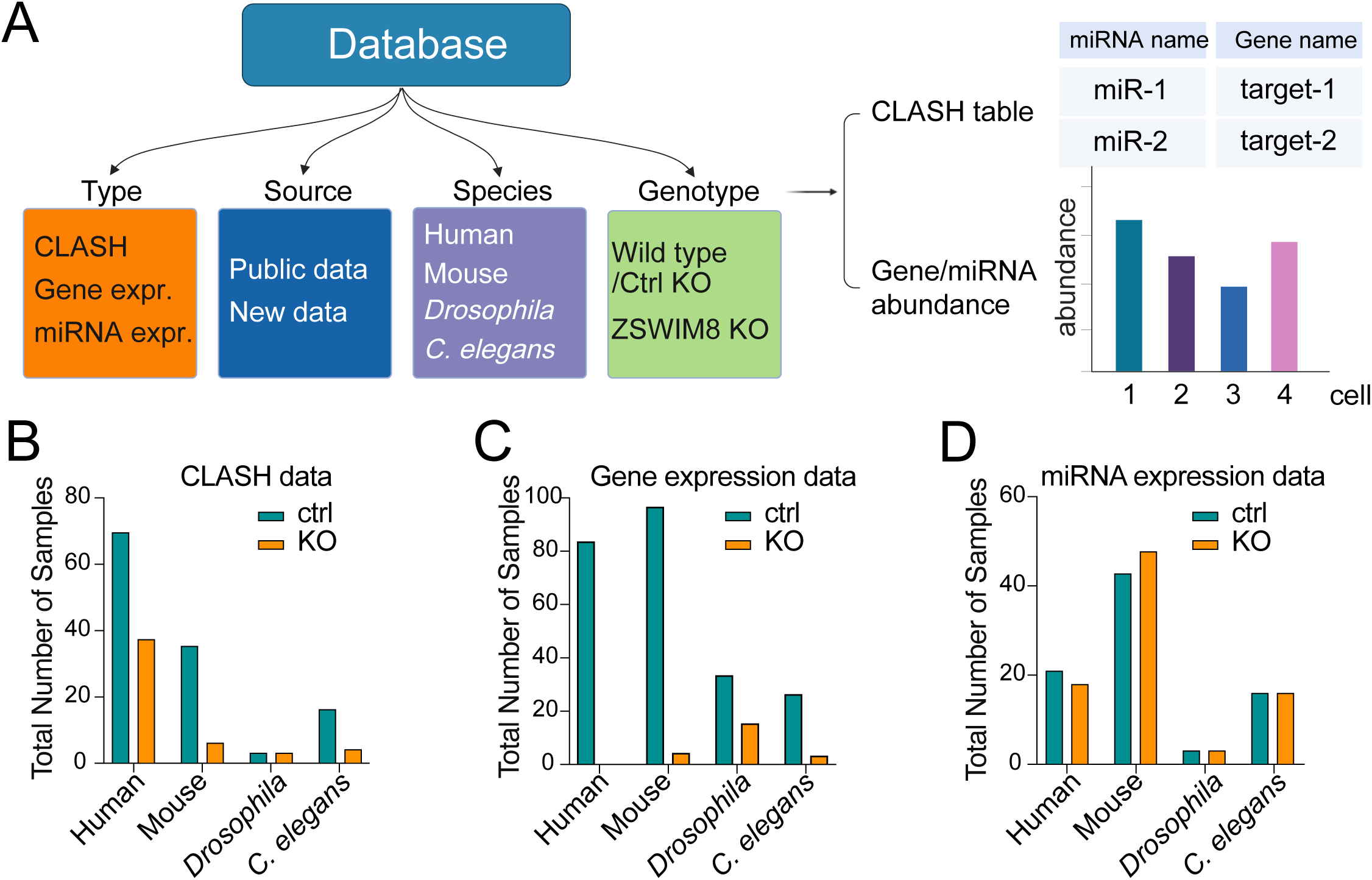
Overview of the datasets integrated in CLASHub. (A) Schematic representation of the CLASHub database composition. The platform integrates three primary data types (orange)—CLASH data, gene expression data, and microRNA expression data—sourced from both publicly available studies and newly generated datasets in this work (blue). Data span four species (purple): human, mouse, *Drosophila*, and *C. elegans*, and include multiple experimental genotypes (green): wild-type, non-targeting controls, and ZSWIM8 knockout samples. The database enables retrieval of CLASH hybrid tables and associated RNA and miRNA abundance information from diverse biological contexts. Bar plots summarizing the number of datasets for CLASH (**B**), gene expression (**C**) and miRNA expression (**D**) across species, with or without knockout of ZSWIM8 orthologs.

### CLASH database

To enable convenient access to high-confidence miRNA-target interactions, CLASHub features a user-friendly web interface that allows users to search CLASH data in five intuitive steps (Figure 2A, #1-#5). Users can select data type (CLASH), followed by species (e.g., Human), cell line or tissue (e.g., A549), and a miRNA name, gene name, or gene ID of interest (e.g., miR- 7-5p). After clicking “Search”, users are directed to a result page that displays all detected hybrids involving the queried miRNA or gene (Figure 2B). The CLASH data in this database were generated using antibodies specific to different AGO proteins: pan-AGO (AGO1-4) in human and mouse, Ago1 in *Drosophila* and ALG-1/ALG-2 in *C. elegans*.

**Figure 2.**
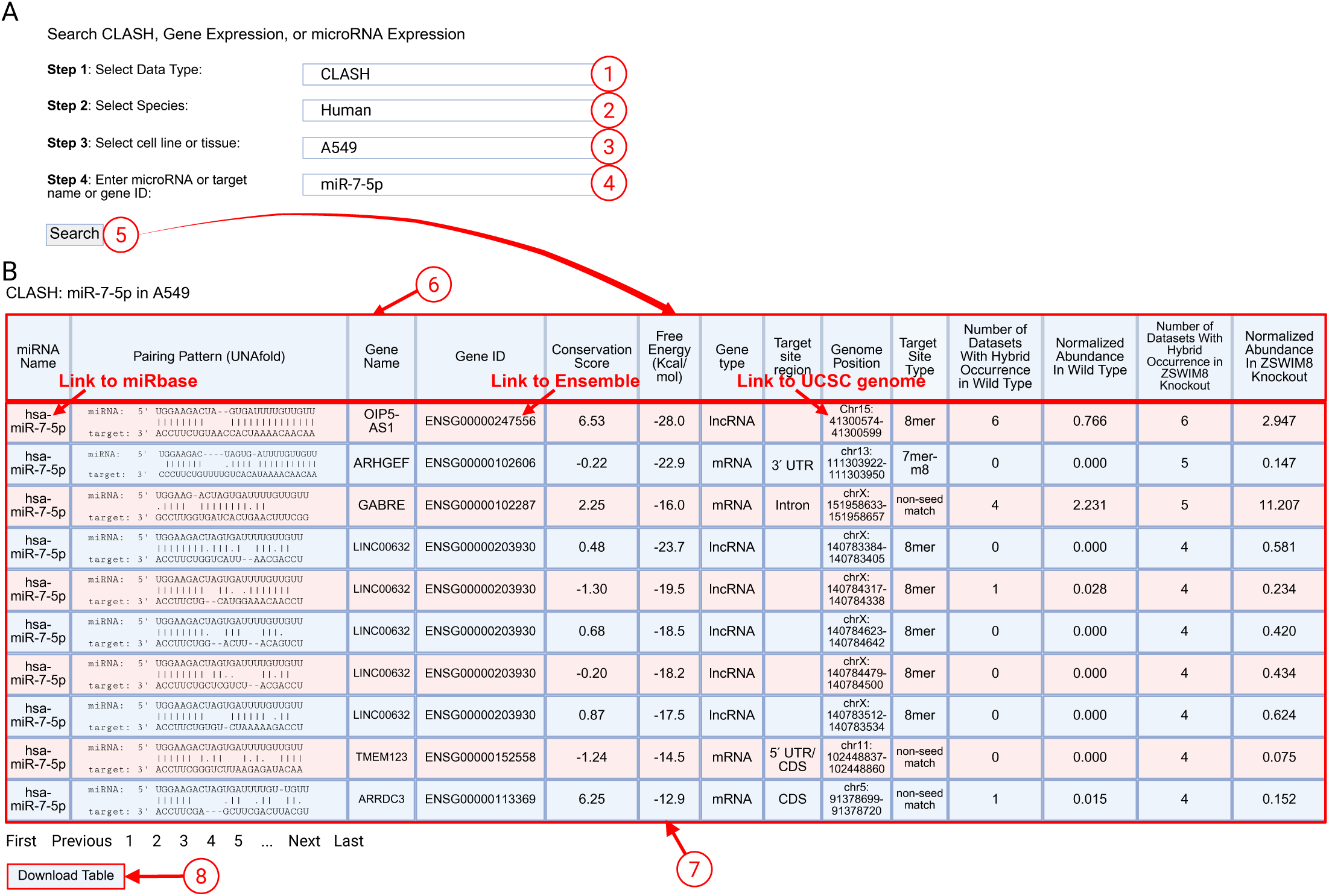
User interface for querying miRNA-target hybrids in the CLASH database. (A) CLASH data could be achieved from the CLASHub web interface guides users by selecting (1) data type, (2) species, (3) sample type and (4) miRNA or gene ID in the query fields, and perform the (5) “Search” function. (B) The example CLASH output table summarizes detected miR-7 hybrids in A549 cells (6). Each row includes comprehensive information regarding the hybrids, such as miRNA name, predicted pairing pattern, etc (7). The “*Download Table”* button allows users to retrieve the data table for offline analysis (8). Interactive links allow users to directly open miRBase (via miRNA name), Ensembl (via gene ID), and UCSC Genome Browser (via genome position) for each hybrid.

As shown in Figure 2B (#6), the result table summarizes each miRNA-target hybrid with key features including the miRNA name, base pairing pattern (predicted by UNAfold (Markham & Zuker, 2008)), target gene name and ID, conservation score, free energy of the predicted duplex, gene type, target site region (e.g., 5′ UTR, CDS, and/or 3′ UTR), genome position, and target site type (e.g., 8mer, 7mer-m8, 7mer-A1, 6mer, offset 6mer, or non-seed match), which reflect different level of binding affinity and repression efficacy. Each hybrid is also annotated with the number of replicates in which it was observed under wild-type (WT) and ZSWIM8-knockout (KO) conditions, along with corresponding normalized abundance values.

To illustrate the platform’s utility, we queried miR-7-5p, a highly conserved and neuron-enriched miRNA implicated in tumor suppression and neural development (Cerda-Jara et al., 2024; Scoyni et al., 2024; Yamada et al., 2023) (Figure 2B, #7). Among the interactions identified in A549 cells, one prominent hybrid involves *OIP5-AS1* (also known as *Cyrano*), a long non-coding RNA previously validated as a TDMD trigger for miR-7 in the mammalian brain (Kleaveland et al., 2018). Loss of the *Cyrano* miR-7 binding site has been shown to dramatically elevate miR-7 levels—by 2- to 40-fold—in both cultured cells and murine neuronal tissues, such as the cerebellum (Kleaveland et al., 2018). In our dataset, this hybrid is reproducibly detected in 6 replicates of both WT and KO conditions, with a clear increase in normalized abundance upon ZSWIM8 knockout (from 0.766 to 2.947, Figure 2B), consistent with increased miRNA abundance after loss of TDMD.

Another notable example is the hybrid between miR-7-5p and *LINC00632*, a locus encompassing ciRS-7, a circular RNA previously described as an efficient miRNA sponge for miR-7 (Hansen et al., 2013). Detection of these interactions demonstrates the ability of CLASHub to reveal non-coding RNA interactions and adds further support to the functional relevance of circular RNAs in miRNA regulation.

Importantly, each row in the results table provides interactive links to external resources, enabling seamless transition to explore detail information about miRNA, gene, and genomic context (Figure 2B, #7). Specifically, clicking on the miRNA name, gene ID, or genome position directs the user to the corresponding entries in miRbase (Figure S2A), Ensembl (Figure S2B), and the UCSC Genome Browser (Figure S2C), respectively. To illustrate these link-out functionalities, Figure S2 shows representative examples of the landing pages when querying hsa-miR-7-5p, ENSG00000247556 (*OIP5-AS1*), and chr15:41300574-41300599 respectively. Users can view the results table in the web browser or download the full dataset with a single click (Figure 2, #8).

### Gene and miRNA expression databases

To facilitate interpretation of miRNA-target interactions, CLASHub integrates expression modules that provide both gene and miRNA abundance profiles across diverse biological contexts, based on RNA-seq and miRNA AQ-seq datasets. This dual capability enables researchers to assess whether a given miRNA-target hybrid occurs in the context of appreciable expression of both the miRNA and its predicted target transcript.

Figure 3 illustrates how users can query gene and miRNA expression data. Users first select the Gene Expression data type and species of interest, then enter a gene name or Ensembl gene ID (e.g., *Sftpc*). Alternatively, users can select miRNA Expression from the Data Type dropdown menu and enter a miRNA name (e.g., mmu-miR-124-3p) (Fig. 3A). Clicking “Search” generates a bar plot displaying transcript or miRNA abundance across tissues and cell types, quantified as transcripts per million (TPM) or counts per million (CPM). For example, querying the surfactant protein C gene shows marked enrichment in lung samples (Figure 3B), and mmu-miR-124-3p is highly enriched in brain and iNeuron samples, with negligible expression in non-neuronal tissues (Figure 3C).

**Figure 3.**
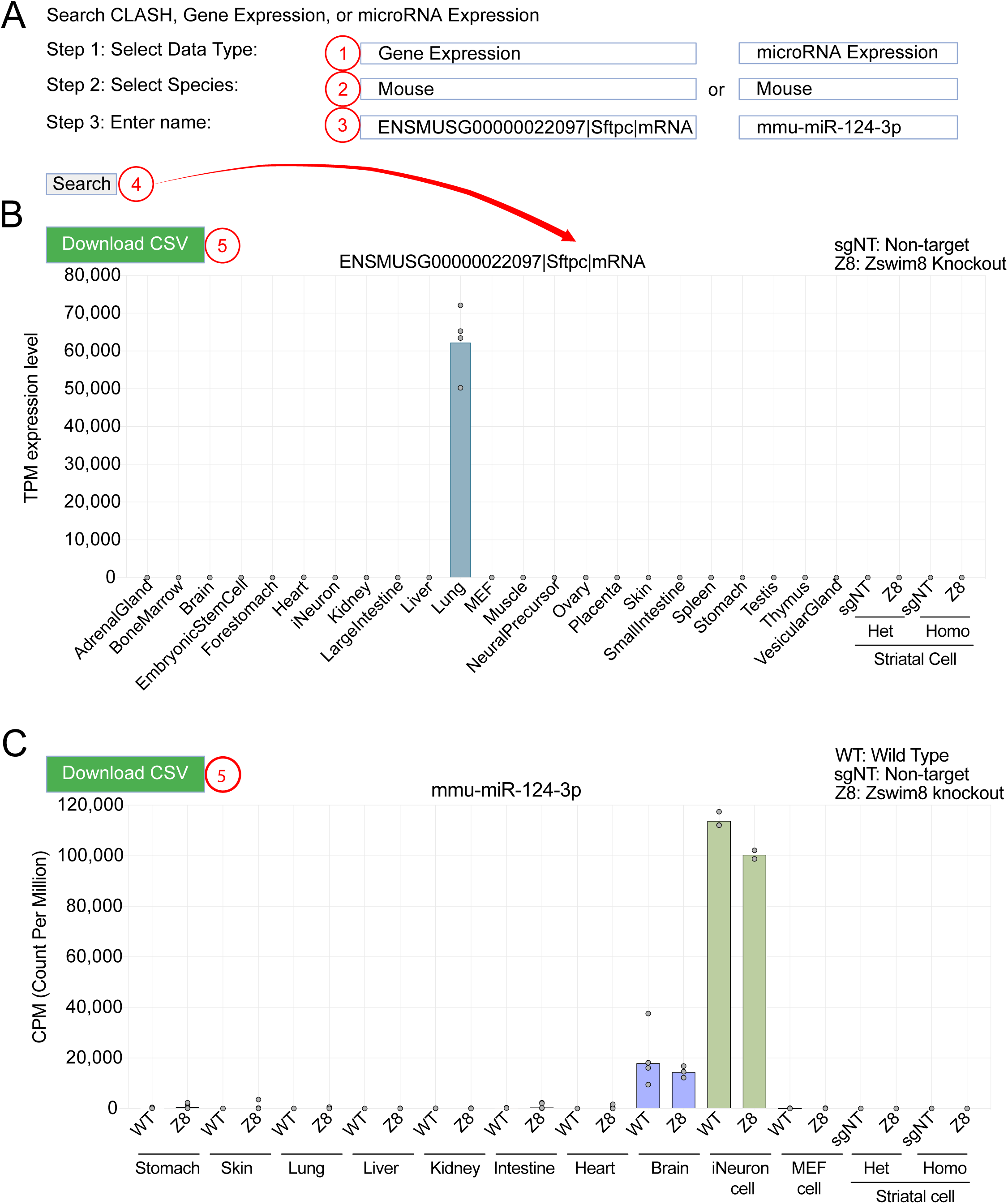
CLASHub gene and miRNA expression database interface and example outputs. (A) User interface for exploring RNA-seq data and miRNA-seq data across tissue and cell types. Users (1) select the “Gene Expression” or “miRNA expression” in the data type, (2) specify the species, and (3) enter a gene name, Ensembl gene ID or miRNA name. (4) Clicking the “Search” retrieves expression profiles. Example output bar plots showing transcript abundance in TPM (B) or miRNA abundance in CPM (C) across mouse tissues and cell types. (5) The “*Download CSV*” button allows exporting the data table for offline analysis.

In both modules, users can export results by clicking the “Download CSV” button (Figure 3B-C, #5). When combined with CLASH hybrid data, these expression profiles provide essential context for evaluating the biological relevance of identified miRNA-target interactions and prioritizing candidates for functional validation.

### CLASHub Analyzers

#### Overview of analytical functionalities provided by CLASHub

Other than summarizing the existing data in databases described above, CLASHub also offers “Analyzer” modules to analyze new CLASH, miRNA-seq, RNA-seq data provided by the users, to generate cumulative fraction curve, and to check job status, all accessible via the left menu (Figure 4). Together, these enable comprehensive miRNA-network studies across the four model species supported in this resource: human, mouse, *Drosophila*, and *C. elegans*. To perform analysis, users can upload raw or pre-processed sequencing data, choose the species, input adapter sequences and output name. The backstage custom scripts will automatically process the data and send results to the users by e-mail. Additionally, the Job Status module allows users to monitor current job counts and query specific job statuses using a unique Job ID, enabling convenient verification of whether a job has completed, is in progress, or remains pending in the queue.

**Figure 4.**
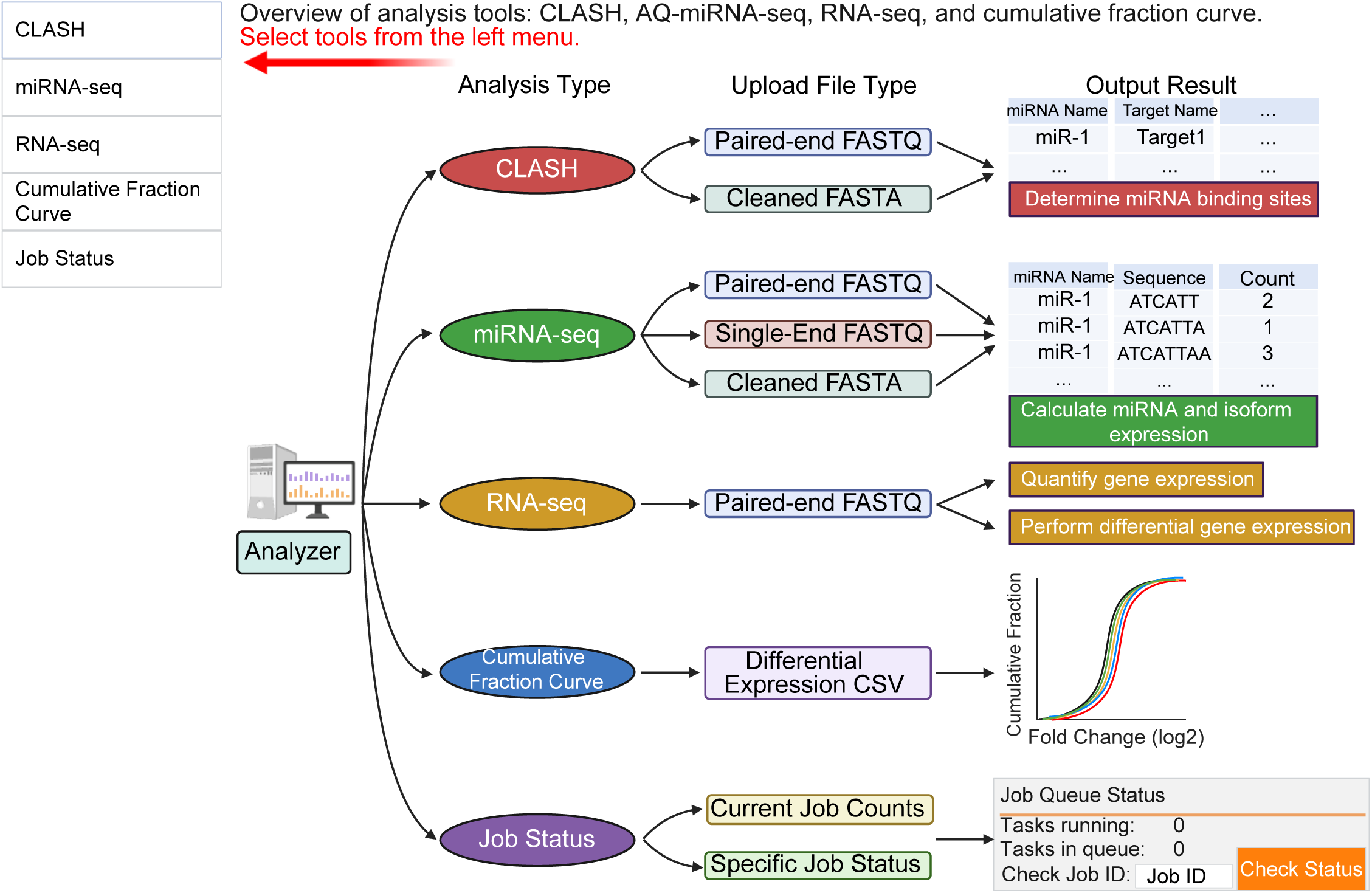
Overview of analysis tools provided by CLASHub. Schematic representation of the five main modules available in the CLASHub Analyzer interface, including CLASH (red), miRNA-seq (green), RNA-seq (orange), cumulative fraction curve generation (blue), and job status monitoring (purple). Users can select the desired analysis type from the left menu. Each module specifies compatible input file formats and outlines the corresponding output content.

All pipelines are fully automated, standardized, and optimized for high-throughput workflows, and all scripts are available at https://github.com/UF-Xie-Lab/CLASHub; Sample input files for testing individual CLASHub Analyzer modules are available (e.g., Figure 5A: Sample CLASH FASTQ 1 and 2). The following sections describe each module in detail.

**Figure 5.**
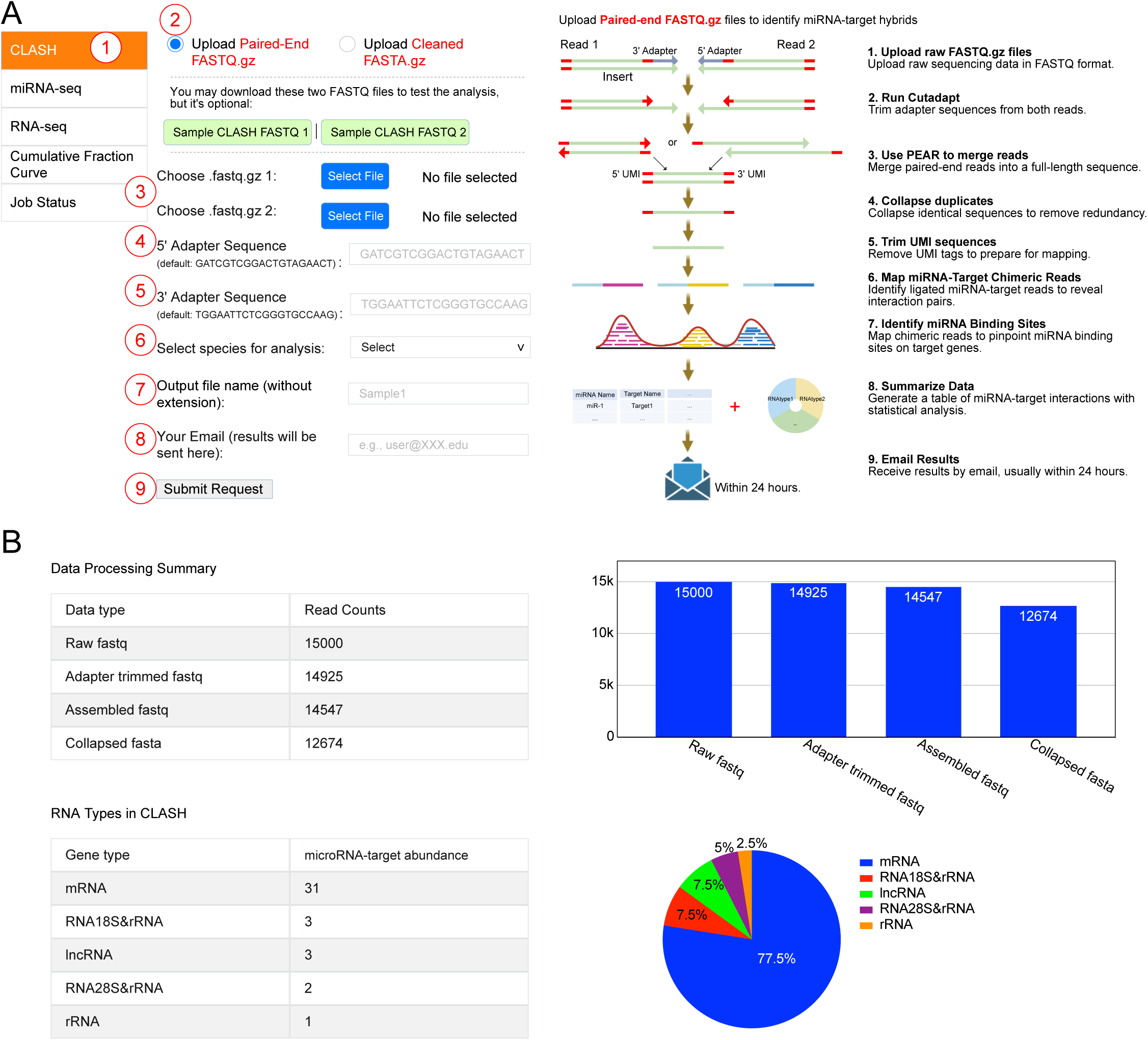
CLASH analysis interface and automated processing pipeline in CLASHub. (A) Interface for initiating CLASH analysis. Users (1) select the CLASH module, (2) select input file type, (3) select local sequencing files for upload, (4 and 5) input adapter sequences, (6) select species, (7) specify output filename, and (8) enter email address, then (9) submit the request. The backstage bioinformatic pipeline to process paired-end FASTQ data is shown on the right. (B) Example output report, displaying processed read counts, RNA type summary, and RNA composition chart.

### CLASH analysis: miRNA-target interaction discovery

The CLASH analysis pipeline in CLASHub identifies miRNA-target chimeric hybrids from raw paired-end FASTQ files or pre-processed single-end FASTA files. For paired-end data, uploaded files undergo sequential processing including adapter trimming (Cutadapt v2.10 (Martin, 2011)), read merging (PEAR v0.9.6 (Zhang et al., 2014)), deduplication (fastx_collapser v0.0.14), and Unique Molecular Identifier (UMI) trimming (Figure 5A). Single-end input files directly proceed to mapping without these initial preprocessing steps (Figure S3).

Chimeric miRNA-target reads are identified using hyb (Travis et al., 2014), aligned to Ensembl reference transcriptomes using Bowtie2 v2.5.3 (Langmead & Salzberg, 2012), and mature miRNA sequences from miRBase Release 22.1. Hybrid free energies (ΔG) are calculated using UNAfold v3.8. miRNA binding sites in the target are annotated with a custom CLASHub.py script, and conservation scores are assessed using phyloP from the UCSC Genome Browser (Perez et al., 2024). Outputs include a detailed hybrid interaction table (miRNA-target pairs, genomic locations, base-pairing patterns, ΔG values, conservation scores, and transcript annotations such as 3ʹ UTR or CDS; Table S4, and an HTML summary report with processing statistics and RNA-type composition (mRNAs, lncRNAs, rRNAs) (Figure 5B). Results are delivered to users by email typically within 24 hours, enabling high-resolution, cross-species exploration of the miRNA-target interaction landscapes.

### miRNA-seq analysis: miRNA quantification and isoform profiling

The miRNA-seq module of CLASHub quantifies miRNA expression and isoform-specific abundances from user-uploaded sequencing data in formats of paired-end FASTQ, single-end FASTQ, or pre-processed cleaned FASTA. Paired-end and single-end FASTQ files (Figure S4 and S5A) undergo preprocessing steps similar to those applied in the CLASH analysis (Figure 5A), which convert the input files to clean FASTA files (Figure S5B).

The CLASHub.py script quantifies miRNA levels by perfectly matching the first 18 nucleotides of each trimmed read to the 5′ end of mature miRNA sequences (miRBase Release 22.1), enabling accurate estimation of total miRNA and isoform abundances. Output includes two tables—one for total miRNA abundance (Table S5) and one for isoform-level counts (Table S6)— as well as a comprehensive HTML report summarizing read processing metrics, filtering statistics, and graphical visualizations of sequence trimming and read distribution (Figure S4B).

### RNA-seq analysis: gene expression quantification and differential expression

CLASHub’s RNA-seq module quantifies gene expression and identifies differentially expressed genes from paired-end FASTQ data. All input reads undergo adapter trimming, genome mapping (HISAT2 (D. Kim et al., 2019)), and quantification of gene expression levels using StringTie (Pertea et al., 2015), producing normalized transcripts per million (TPM) (Figure S6A). Output from this workflow is a gene expression table listing TPM-normalized abundances across samples (Table S7).

For differential expression analysis, users upload paired-end FASTQ files from “Control” and “Treatment” groups. Reads undergo adapter trimming and alignment (HISAT2). Raw gene counts extracted via prepDE.py3 (StringTie developers) are analyzed with DESeq2 (Love et al., 2014), identifying significantly altered gene expression (Figure S7). Results are provided as a differential expression results table containing gene names, log2 fold changes, and adjusted p- values (Table S8), alongside comprehensive HTML reports detailing analysis summaries and visualizations (Figure S6B).

### Cumulative fraction curve generator: functional validation of miRNA impact

The cumulative fraction curve generator evaluates whether miRNA expression changes affect target gene expressions relative to non-targets. Users upload differential gene expression CSV files, which could be generated via the RNA-seq differential expression pipeline in the RNA-seq analysis module, containing essential columns: GeneName, BaseMean (default threshold: 100), and log2FoldChange. Inputs also include species, miRNA name, output file name, and email address (Figure 6). This module simultaneously compares the distribution of expression changes across multiple target types (CLASH-identified targets, Targetscan-predicted targets, etc.), which are displayed together in a cumulative fraction curve plot. This combined display of CLASH- validated and TargetScan-predicted targets within the same plot robustly supports interpretation of miRNA-mediated regulation by allowing direct comparison of their expression against non-target transcripts.

**Figure 6.**
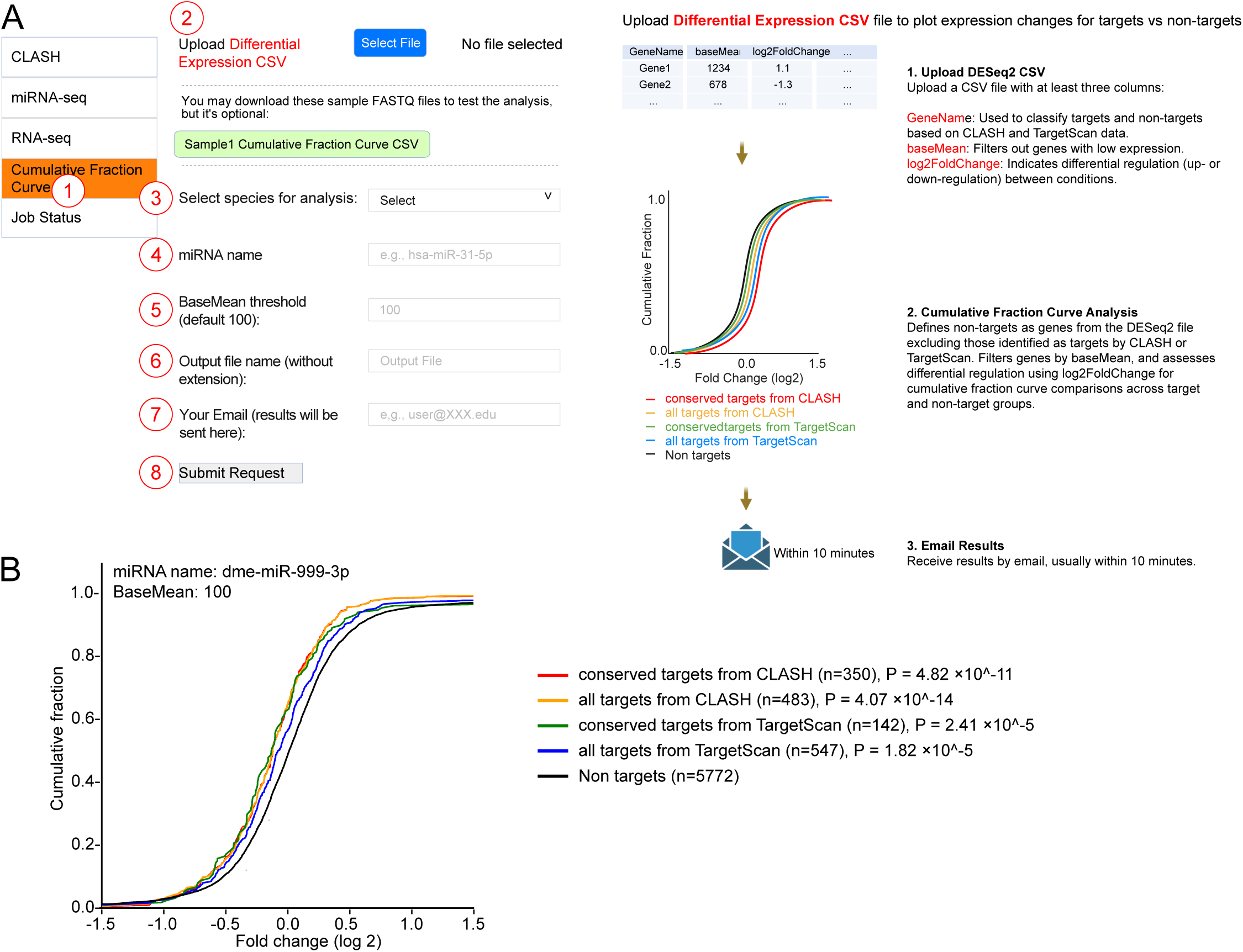
Cumulative fraction curve analysis module and job monitoring interface. (A) Interface for generating cumulative fraction curves. Users (1) select the module, (2) upload a differential gene expression CSV file, (3) specify species, (4) enter miRNA name, (5) define optional miRNA count BaseMean threshold, (6) specify output filename, (7) provide email, and (8) submit the request. The backstage bioinformatic pipeline to generate the plot is shown on the right. (B) Example output showing cumulative fraction curves for target transcripts fold-change with differential expression of dme-miR-999-3p, with Mann–Whitney U test p-values indicating significance of expression shifts in predicted target sets compared to non-targets.

To illustrate this functionality, Figure 6B shows an example of cumulative fraction curves for dme-miR-999-3p target genes. This plot shows the overall reduction in miR-999 target expression in cells with high levels of miR-999 compared to the control condition (Sheng et al., 2023). The module automatically performs Mann–Whitney U tests to quantify statistical differences in expression shifts between each target group and non-target genes. In addition to graphical visualization, the output also includes a downloadable table (Table S9) that annotates each gene in the user-provided differential expression file with its target classification (e.g., conserved CLASH target, conserved TargetScan target, all CLASH targets, all TargetScan targets, or non-target). This enables an easy review of how each gene contributes to the observed distributions and facilitates downstream analyses or reporting.

### Insights into miR-335-3p regulation by TDMD using CLASHub

To demonstrate the integrated capabilities of CLASHub as both a curated miRNA-target interaction database and an analysis platform, we applied it to explore miR-335-3p, a miRNA known to be regulated by TDMD (Shi et al., 2023; Jones et al., 2023). Prior studies have consistently shown that miR-335-3p is among the most strongly upregulated miRNAs when ZSWIM8 is knocked out, indicating its potential regulation by TDMD. Motivated by these findings, we leveraged both the CLASHub database and analysis modules to investigate miR-335-3p expression, potential TDMD triggers, and binding site conservation across species.

Using CLASHub’s miRNA expression datasets, miR-335-3p expression was analyzed across samples from human cell lines to mouse tissues. In ZSWIM8-knockout human cells (e.g., HeLa and A549) and Zswim8-knockout mouse samples, which include cells such as MEF and tissues like the brain, kidney, lung, and stomach, miR-335-3p expression was 3- to 12-fold higher compared to wild-type or non-targeting knockout controls (Figure 7A). The consistent response across diverse human and mouse samples highlights miR-335-3p’s universal sensitivity to ZSWIM8/Zswim8 depletion.

**Figure 7.**
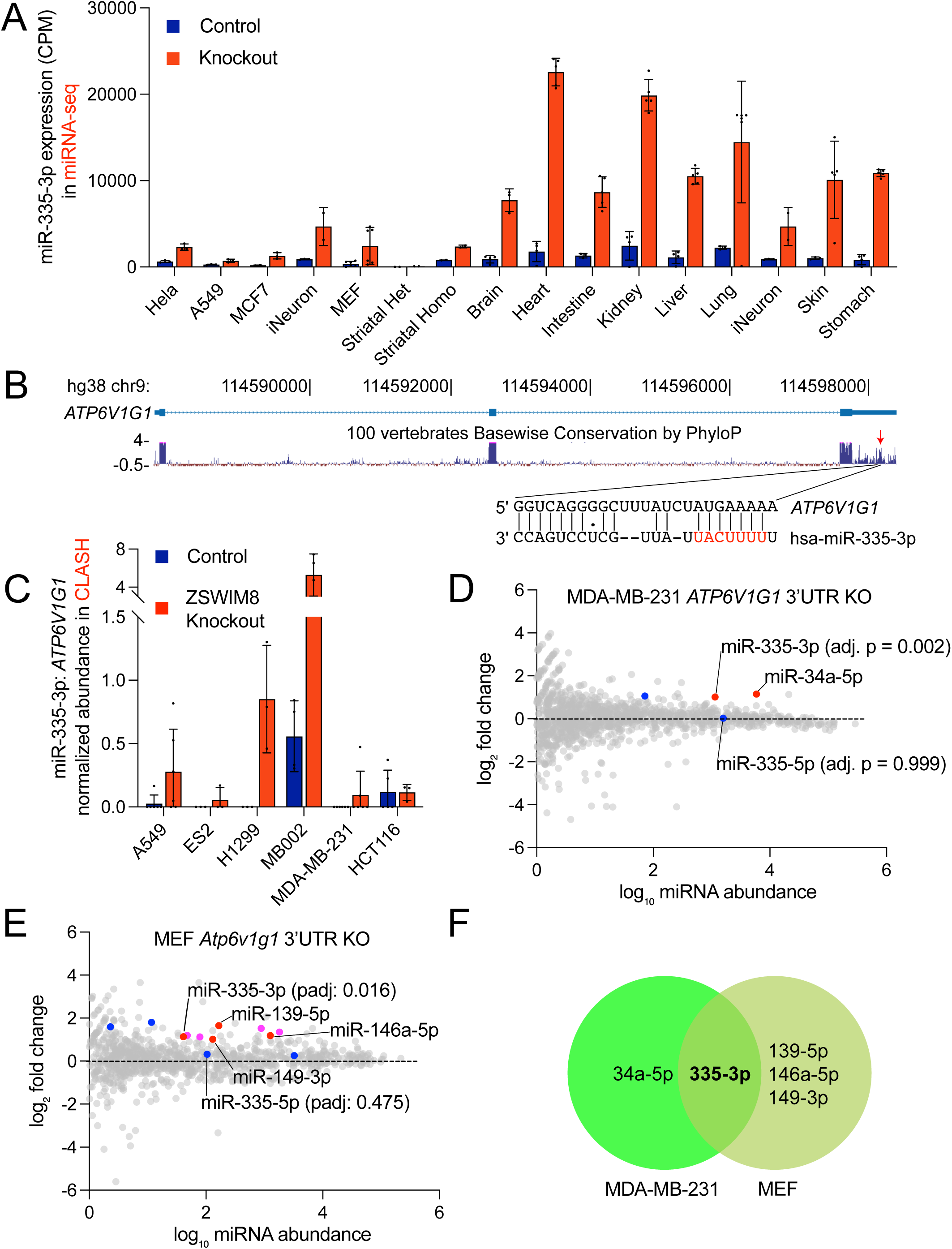
Insights into miR-335-3p Regulation and Trigger Discovery Using CLASHub. (A) miR-335-3p levels in ZSWIM8-knockout (red) and control (blue) samples across human cell lines and mouse tissues. Expression values are shown as CPM from miRNA AQ-seq datasets. (B) UCSC Genome Browser view of the *ATP6V1G1* 3′ UTR on chromosome 9. The miR-335-3p binding site was denoted with a red arrow. PhyloP conservation track highlight conserved regions with blue bars. (C) Normalized abundance of miR-335-3p:*ATP6V1G1* hybrids in CLASH datasets across six human cell lines. (D-E) Differential expression of mature miRNAs in (D) MDA-MB-231 *ATP6V1G1* 3′UTR knockout and (E) MEF *Atp6v1g1* 3′UTR knockout compared to control cells. Each point represents a mature miRNA, plotted by log10 abundance (x-axis) and log2 fold change (y-axis). miRNAs significantly upregulated in knockout samples (adjusted p-value <0.02) are shown in red. Corresponding passenger strands are indicated in blue if not significantly changed. (F) Venn diagram showing overlap of significantly upregulated miRNAs between MDA-MB-231 and MEF cells, upon *ATP6V1G1*/*Atp6v1g1* trigger KO.

To pinpoint potential triggers driving miR-335-3p TDMD, we focused on identifying potential miRNA-target interactions. Our analysis with CLASHub identified over two hundred miR-335-3p hybrids across six human cell lines under both control and ZSWIM8-knockout conditions, including A549, ES2, H1299, HCT116, MB002, and MDA-MB-231, where miR-335- 3p expression is strongly upregulated, facilitating robust hybrid detection and comparative analysis.

To prioritize the most likely TDMD candidates, we applied stringent criteria requiring perfect seed pairing at positions 2-8 of the miRNA and at least eight contiguous nucleotides of complementarity within the miRNA 3ʹ end (Buhagiar & Kleaveland, 2024; Li et al., 2021). This filtering strategy yielded seven high-confidence miR-335-3p hybrids distributed across the six cell lines (Table 1).

**Table 1.**
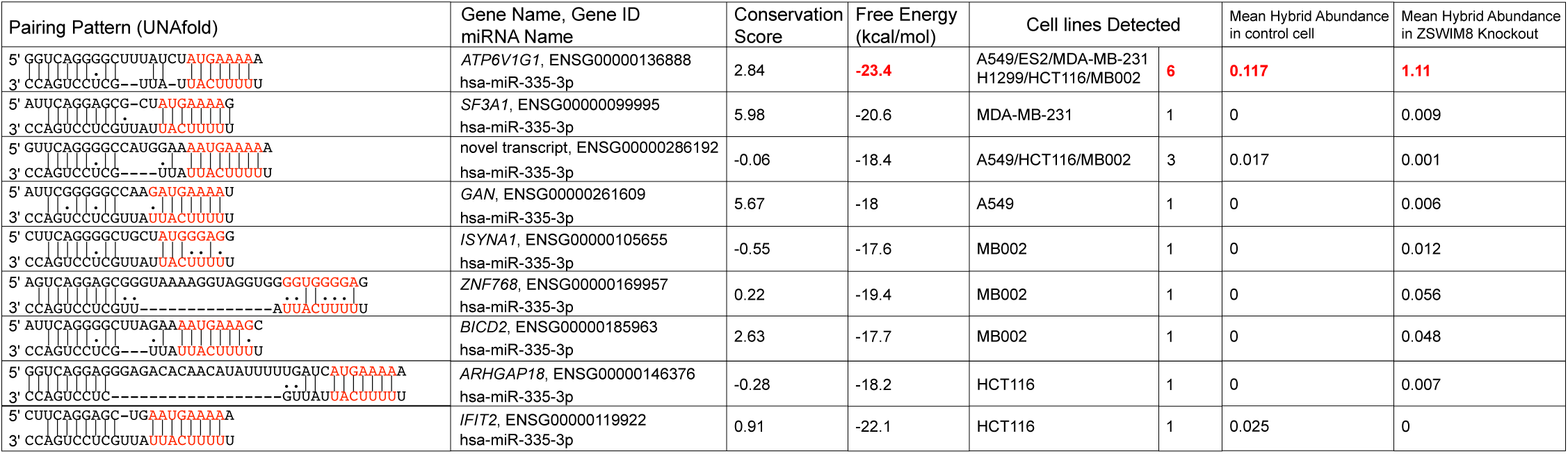
Candidate miR-335-3p TDMD Hybrids Detected in Six Cell Lines.

Among all miR-335-3p hybrids detected, the interaction with *ATP6V1G1* stood out due to several features: First, its base-pairing pattern includes a contiguous 7-nucleotide seed match and 10 consecutive nucleotides of complementarity at the miRNA 3ʹ end, a configuration characteristic of TDMD-inducing sites (Figure 7B and S8A). Second, it exhibited a highly favorable predicted free energy of –23.4 kcal/mol, indicating strong thermodynamic stability. Third, *ATP6V1G1* hybrids were detected consistently across all six cell lines, whereas the other candidate hybrids were observed in only one or two cell lines, reflecting limited reproducibility across samples. Fourth, the mean hybrid abundance was markedly higher both in control cells (0.117) and ZSWIM8 knockout cells (1.314) compared to the other hybrids (Figure 7C and Table 1). Finally, PhyloP conservation analysis across vertebrates demonstrated significant conservation at the seed and 3′ end pairing regions. Notably, this miR-335-3p:*ATP6V1G1* interaction has not been previously reported, further supporting its potential relevance as a novel TDMD trigger.

To experimentally validate *ATP6V1G1*’s involvement in miR-335-3p TDMD, we employed CRISPR/Cas9 to remove the potential TDMD trigger sequence. Three homozygous knockout clones (M1: 108 bp deletion with 1 bp insertion, M5: 118 bp deletion with 6 bp insertion, and M8: 108 bp deletion) were generated in human MDA-MB-231 cells (Figure S8B), and four homozygous knockout clones (b5, b6, b11, and b17, all with identical 65 bp deletions) were generated in mouse MEF cells (Figure S8C).

To reduce false positives in low-abundance miRNAs, we applied stringent thresholds (baseMean >40, adjusted p-value <0.02, log2 fold change >1) when analyzing the miRNA-seq data in *ATP6V1G1/Atp6v1g1* trigger knockout cells. Specifically, in MDA-MB-231 cells, miR- 335-3p and miR-34a-5p were the most prominently elevated (Figure 7D, red dots), while their corresponding passenger strands were not significant upregulated (Figure 7D, blue dots). In MEF cells, miR-335-3p, miR-139-5p, miR-146a-5p, and miR-149-3p showed marked increases (Figure 7E, red dots), with their passenger strands not significantly upregulated (Figure 7E, blue dots). Notably, miR-195a and miR-497 exhibited concurrent upregulation of both 5p and 3p strands, suggesting that their apparent changes may be due to increased transcription rather than loss of TDMD (Figure 7E, magenta dots). Overall, miR-335-3p was only miRNA upregulated in both human and mouse trigger knockout models, reinforcing the specific role of *ATP6V1G1/Atp6v1g1* as a TDMD trigger (Figure 7F, Table S10-11).

## Discussion

Multiple high-throughput experimental methods, such as HiTS-CLIP, PAR-CLIP, CLASH, and Chimeric eCLIP, have been developed to identify miRNA binding sites (Anastasakis et al., 2021; Gay et al., 2018; Manakov et al., 2022; O’Connor et al., 2021). However, existing studies are often limited to a narrow range of cell lines and tissues, including human cell lines such as HCT116, HEK293T, Huh-7.5, TIVE-EX-LTC, as well as mouse cell lines like HE2.1B, 3T12 and cortex tissues, among others (Whitney L Bullard et al., 2019; W. L. Bullard et al., 2019; Fields et al., 2021; Gay et al., 2018; Manakov et al., 2022; Moore et al., 2015). In this study, beyond the incorporation of publicly available CLASH data, additional datasets from a diverse range of cancer cell lines and tissues were utilized or generated, including human cell lines (e.g., A549, D425, ES2, HepG2, H1299, MB002, MDA-MB-231, OVCAR8, T98G, U87MG, and 501Mel), mouse samples (e.g., MEF, striatal cells, heart tissue, and kidney tissue). The expanded scope of CLASH data includes both control samples (e.g., wild type or non-targeting sgRNA) and ZSWIM8-knockout samples, which are essential for studying TDMD (Grimme et al., 2025; Han et al., 2020; Jones et al., 2023; LaVigne et al., 2025; Li et al., 2021; Shi et al., 2023; Stubna et al., 2025). Furthermore, gene expression and miRNA expression data from publicly available datasets have been re- analyzed and integrated into the CLASHub database. These datasets provide valuable insights and enable researchers to efficiently explore miRNA-target interactions across various experimental conditions.

In addition to its database functionality, CLASHub offers an “Analyzer” interface, a graphical user interface that simplifies miRNA-related data analysis. Unlike previous tools, such as Hyb, which identifies miRNA-target hybrids from CLASH FASTQ files but lacks advanced data processing (Travis et al., 2014), the CLASHub analyzer automates the entire analytical workflow. A recently developed Hybkit provides a powerful Python-based toolkit for hybrid data analysis, but it requires coding skills and is less accessible to non-programmers (Stribling et al., 2023). In contrast, CLASHub allows users to upload their own FASTQ or FASTA files, and the platform processes the data, providing results via email. These results include miRNA-target hybrid abundances, miRNA binding site conservation scores, and miRNA-target pairing patterns. Beyond CLASH analysis, the platform also supports miRNA-seq and RNA-seq analyses, as well as cumulative fraction curve analyses based on miRNA-regulated differential gene expression. By offering these integrated features, CLASHub significantly reduces the time and computational expertise required for miRNA-related analyses, allowing researchers to focus on biological interpretation.

### CLASHub identifies *ATP6V1G1* as a TDMD trigger of miR-335-3p

miR-335-3p has been previously reported to play critical roles in memory function in the brain (Raihan et al., 2018). Additionally, it functions as a tumor suppressor in several cancers and targets LRRK2 to reduce inflammation, potentially mitigating Parkinson’s disease progression (Oliveira et al., 2021). These findings underscore the significant roles of miR-335-3p in neurological and cancer-related pathways. Furthermore, prior studies have demonstrated that miR- 335-3p is regulated by ZSWIM8-mediated TDMD in various mouse tissues (Jones et al., 2023); however, the specific trigger responsible for miR-335-3p degradation has remained unclear.

Using the CLASHub platform, a potential trigger for miR-335-3p degradation was identified in the 3ʹ UTR of *ATP6V1G1*. This finding was further validated through the CLASHub platform’s Analyzer function, where miR-seq analysis was employed to assess miRNA abundance changes in *ATP6V1G1* trigger knockout samples. These analyses confirmed that the identified *ATP6V1G1* trigger plays a role in miR-335-3p degradation. Notably, miR-335-3p abundance increased approximately two-fold upon *ATP6V1G1/Atp6v1g1* deletion in MDA-MB-231 and MEF cells (Figure 7D-E), whereas ZSWIM8 knockout in MEF cells resulted in a ∼7-fold upregulation (Figure 7A), suggesting that additional TDMD triggers likely contribute to miR-335-3p turnover. This discovery not only provides new insights into miRNA regulatory mechanisms but also highlights the capability of CLASHub to uncover and characterize critical miRNA-target interactions.

The development of CLASHub represents a significant advance in simplifying miRNA research workflows. It facilitates both exploratory and in-depth analyses, addressing limitations in existing tools and providing a more comprehensive platform for researchers. This study demonstrates the potential of CLASHub to serve as a steppingstone toward a deeper understanding of miRNA landscapes in various biological contexts. To further enhance CLASHub’s utility, community feedback and suggestions for new features are welcomed to guide its continuous development.

## Materials and Methods

### CLASH Experiment and Analysis

CLASH experiments performed in this study followed protocols adapted from qCLASH (Fields et al., 2021; Sheng et al., 2023) or chimeric eCLIP (Manakov et al., 2022), as specified in Table S1. Briefly, cells were UV crosslinked on ice at 254 nm with a total energy of 400 mJ/cm² to covalently link RNA–protein complexes.

For qCLASH, ∼200 µL cell pellet was lysed in 1 mL cold lysis buffer on ice for 15 minutes, followed by incubation at 37°C for 5 minutes with shaking at 1000 rpm. Lysates were centrifuged at 21,000 g for 15 minutes at 4°C. Subsequently, 3 mg of protein lysate underwent overnight immunoprecipitation at 4°C using either 40 µL of Protein L beads (Thermo Fisher, PI88850) conjugated with approximately 80 µg anti-AGO2 4F9 antibody (human and mouse samples) or 200 µL of Protein A beads (Life Technologies, #10002D) pre-bound with 20 µg anti-Drosophila Ago1 antibody (Abcam, ab5070). Immunocomplexes were washed three times with 950 µL lysis buffer, treated with 150 µL of 15 ng/mL RNase A for 12 minutes at 22°C, and sequentially washed three times with 950 µL of each of the following buffers: 1X PXL, 5X PXL, High Stringency Buffer, High Salt Buffer, and PNK buffer (buffer compositions and reaction mixes are listed in Table S12). RNAs on beads were phosphorylated with 80 µL T4 PNK mix at 16°C for 40 minutes, washed three times with 950 µL PNK buffer, and subjected to intermolecular ligation using 500 µL Intermolecular Ligation Mix overnight at 4°C. Following ligation, beads underwent three washes with 950 µL 1X PNK buffer, were treated with 80 µL dephosphorylation mix at 16°C for 40 minutes with intermittent shaking (15 s every 1 min at 1200 rpm), washed twice with 950 µL 1X PNK-EGTA and three times with 950 µL 1X PNK buffer, and incubated overnight at 16°C in 80 µL 3′-adapter ligation mix with intermittent shaking (15 s every 1 min at 1400 rpm) (adaptor sequences are listed in Table S12). After washing three times with 950 µL 1X PNK buffer, protein complexes were eluted by incubation with 100 µL protein elution buffer at room temperature for 15 minutes with continuous shaking at 1400 rpm. Eluted AGO complexes were digested with 50 µL proteinase K mix at 37°C for 20 minutes. After reaction, RNA was extracted by Phenol:Chloroform:Isoamyl Alcohol (PCA) with ethanol precipitation, phosphorylated with 4.5 µL T4 PNK mix at 16°C for 40 minutes. The reaction was extracted again with PCA and precipitated by ethanol, and resuspended into 10 µL H₂O. For 5′ adapter ligation, RNA was mixed with 10 µL of 5′ adapter ligation mix overnight at 16°C. RNA was then purified with PCA and precipitated with ethanol, resuspended in 11 µL H₂O, mixed with 1 µL 10 µM RTP and 1 µL 10 µM dNTPs, incubated at 65°C for 5 minutes, and subjected to reverse transcription using RT mix.

For chimeric eCLIP, cell lysates were prepared in 1 mL ice-cold iCLIP lysis buffer supplemented with 20 μL Murine RNase Inhibitor (NEB, M0314) (buffer compositions and reaction mixes are listed in Table S13). Samples were sonicated using a Bioruptor (low setting, 4°C, 5 min, 30 s on/off cycles), treated with 10 μL Turbo DNase (Invitrogen, AM2239) and 20 μL 1:100 diluted RNase I, and incubated at 37°C for 5 minutes at 1200 rpm. Lysates were cleared by centrifugation (21,000 g, 10 min, 4°C). Immunoprecipitation was performed with 40 µL Protein L magnetic beads conjugated with 80 µg anti-AGO2 4F9 antibody (human and mouse), incubated overnight at 4°C with rotation. Beads were washed thrice each with 500 µL cold High Salt Wash Buffer and Wash Buffer, then once with 200 µL cold 1X PNK buffer. Phosphorylation on beads used 100 µL T4 PNK Minus Mix at 37°C for 20 minutes, followed by one wash with 500 µL High Salt Wash Buffer and three washes with 500 µL Wash Buffer. Intermolecular ligation was conducted overnight at 4°C using 180 µL Intermolecular Ligation Mix. Beads were then washed three times each with 500 µL cold High Salt Wash Buffer and Wash Buffer. FastAP treatment was performed at 37°C for 10 minutes with 50 µL FastAP mix. To the beads with FastAP mix, 150 µL PNK mix was added, followed by 20 minutes incubation at 37°C. This mixture was removed from the beads, followed by one wash with 500 µL High Salt Wash Buffer and three washes with 500 µL Wash Buffer. For 3′ linker ligation, beads were resuspended in 80 µL 3′ adapter ligation mix and incubated overnight at 16°C with intermittent shaking (15 s every 1 min, 1400 rpm). After ligation, four washes followed (once 500 µL cold Wash Buffer, once 500 µL High Salt Wash Buffer, and twice with 500 µL Wash Buffer). Samples were denatured in 7.5 µL 4X LDS buffer (ThermoFisher, NP0008), 3 µL 1 M DTT, and 20 µL Wash Buffer at 70°C for 10 minutes, separated on NuPAGE gels, transferred to nitrocellulose membranes (Amersham Protran 0.45 µm), and a region from 100–250 kDa was excised and digested with 150 µL Proteinase K SDS mix at 37°C for 20 minutes, followed by an additional 20-minute incubation at 50°C. RNA was recovered by PCA extraction and ethanol precipitation and subsequently resuspended in 9 µL nuclease-free water. For reverse transcription, 1 µL of 10 mM dNTPs and 0.5 µL of RTP primer were added to the RNA, incubating the mixture at 65°C for 3 minutes, then immediately placing it on ice. After adding 10 µL RT mix, the reaction was incubated at 55°C for 20 minutes. Following reverse transcription, cDNA was treated with 2.5 µL ExoSAP-IT (ThermoFisher, 78201.1) at 37°C for 15 minutes. The reaction was subsequently mixed with 1 µL 0.5 M EDTA, treated with 3 µL 1 M NaOH at 70°C for 10 minutes, and neutralized using 3 µL 1 M HCl. The cDNA was then purified by binding to 5 µL Silane beads (Invitrogen, 37002D) in the presence of 90 µL RLTW buffer and 108 µL 100% ethanol at room temperature for 10 minutes. The beads underwent two washes with 300 µL of 80% ethanol, followed by a final wash with 150 µL of 80% ethanol. Beads were air- dried for 5 minutes prior to the subsequent ligation step. For the 5′ adapter ligation, cDNA was initially denatured with 1.45 µL TT Elution Buffer, 0.5 µL RA5 DNA adapter (100 µM), and 0.8 µL DMSO at 70°C for 2 minutes. Following denaturation, 7.55 µL of 5′ adapter ligation mix was added, and reactions were incubated overnight at room temperature with rotation. Post-ligation, cDNA was purified via a second silane bead cleanup (using 2.5 µL beads with 45 µL RLTW buffer and 45 µL ethanol), washed as described previously, and eluted into 22 µL Bead Elution Buffer.

For both qCLASH and chimeric eCLIP methods, cDNA libraries were PCR-amplified for 12 cycles using RPI1 and RPIX primers. Amplified libraries were size-selected on 8% polyacrylamide gels to enrich fragments between 180 bp and 400 bp. Sequencing was performed using Illumina NovaSeq platforms. Detailed reagent compositions and oligonucleotide sequences for qCLASH and chimeric eCLIP are provided in Table S12 and Table S13, respectively.

### Identification of miR-335-3p TDMD Triggers

To identify candidate TDMD triggers, all miR-335-3p–target hybrid reads were retrieved from the CLASHub database across six human cell lines (A549, ES2, H1299, HCT116, MB002, and MDA-MB-231) under both control and ZSWIM8-knockout conditions. Hybrids were filtered based on two criteria: perfect base pairing within the seed region (positions 2–8), and at least eight contiguous base pairs within the miRNA 3′ region, with both regions allowing G:U wobble pairing. All analyses were performed using custom scripts available at https://github.com/UF-Xie-Lab/CLASHub.

### Small RNA Sequencing and Analysis

Small RNA libraries were prepared using a modified AQ-seq protocol (Kim et al., 2019). Total RNA was extracted from MDA-MB-231 and MEF cells (either non-targeting sgRNA controls or *ATP6V1G1*/*Atp6v1g1* TDMD region knockouts) using TRIzol reagent (Thermo Fisher Scientific), followed by isopropanol precipitation. For each sample, 15 μg of total RNA was mixed with 1 μL of 3.33 nM synthetic spike-in RNA pool containing 30 distinct sequences, and the mixture was resolved on 15% urea–polyacrylamide gel. RNA fragments of ∼18–30 nt were excised, eluted overnight in 800 μL of 0.3 M NaCl at 25 °C with shaking (1400 rpm), and ethanol precipitated by adding 40 μL 3 M NaOAc (pH 5.2), 0.5 μL GlycoBlue, and 1 mL pre-chilled 100% ethanol. RNAs were resuspended in 3 μL RNase-free water. For 3′ adapter ligation, 3 μL of purified RNA was combined with 0.5 μL of 5 μM miRCat-33 3′ adapter, heated at 70 °C for 2 minutes, and transferred to ice. Then, 6.5 μL of 3′ ligation mix was added, and the reaction was incubated at 25 °C for 4.5 hours. Ligation products (∼45–57 nt) were size-selected on 15% urea– PAGE and eluted overnight in 730 μL 0.3 M NaCl. The eluted RNA was ethanol-precipitated as above and resuspended in 2.5 μL of 5′ adapter mastermix (2.14 μL RNA, 0.36 μL of 5 μM 5′ adapter). The mixture was denatured at 70 °C for 2 minutes and chilled on ice. Subsequently, 7.5 μL of 5′ ligation mix was added, followed by a 1-hour incubation at 37 °C. For reverse transcription, 10 μL of the 5′ and 3′ adapter-ligated RNA was mixed with 1 μL of 4 μM RT primer (RTP), heated at 70 °C for 2 minutes, and placed on ice. A 9 μL RT mix was then added. The reaction was incubated at 50 °C for 1 hour, then heat-inactivated at 70 °C for 15 minutes, and held at 4 °C. cDNA was amplified using RP1 and RPIX primers with 12 PCR cycles. PCR products were resolved on 8% native PAGE, and miRNA insert-sized fragments (∼145 bp) were gel-excised, eluted in 0.3 M NaCl overnight, and ethanol-precipitated. Raw reads were processed and analyzed using the CLASHub Analyzer module. Reagents, buffers, adapters, and spike-in sequences used in small RNA sequencing are listed in Table S14.

### miRNA targets identification from CLASH and TargetScan

CLASH-derived targets require a seed match (8mer, 7mer or 6mer) and are limited to transcripts annotated as either mRNA or non-coding RNA (ncRNA), excluding rRNAs, tRNA, snRNAs, snoRNAs, and all pseudogenes. To be classified as “Conserved targets”, human and mouse interaction must be detected in CLASH hybrid reads across at least two cell types, contain 8mer or 7mer seed matches, have a phyloP conservation scores > 0, and be annotated as mRNA/ncRNA. To determine conservation score, we used UCSC Genome Browser phloP: g38.phyloP100way (human), mm39.phyloP35way (mouse), dm6.phyloP124way (*Drosophila*), and ce11.phyloP135way (*C. elegans*). For *C. elegans*, targets detected at any developmental stage (embryo, L3, or L4) are considered conserved, without requiring recurrence. For *Drosophila*, where only S2 cell data exist, no recurrence criterion is applied. In parallel, TargetScan-predicted targets are extracted from "Summary_Counts.txt" files (Human/Mouse Release 8; *Drosophila* Release 7.2; *C. elegans* Release 6.2), including conserved and non-conserved sites.

### Implementation and Accessibility

CLASHub is a custom web application built with HTML, PHP, JavaScript, and CSS, and sbatch scripts. Hosted on the University of Florida’s HiPerGator infrastructure (PubApps), it leverages scalable computational and database resources to provide interactive visualization, analysis, and download of microRNA-related data for high-performance research. The source code for the web application, including sbatch scripts, PHP modules, and other implementation details, is available at https://github.com/UF-Xie-Lab/CLASHub.

## Data availability

Newly generated sequencing datasets from this study are available in the NCBI SRA under accession number PRJNA1166120. Information for all previously published datasets used in CLASHub—including CLASH, gene expression, and miRNA expression—is summarized in Supplementary Tables S1–S3, and additional details available at https://clashub.rc.ufl.edu/more_info.html. Sample input files for testing the CLASHub Analyzer modules are available at https://clashub.rc.ufl.edu/Analyzer.html (e.g., Figure 5A: Sample CLASH FASTQ 1 and 2). Previously published CLASH datasets include those from PRJNA1093144, GSE303817, PRJNA896239, GSE198250, GSE164634, GSE124687, GSE101978, GSE73057, GSE73058, GSE56180, PRJNA328816 (Broughton et al., 2016; W. L. Bullard et al., 2019; Fields et al., 2021; Gay et al., 2018; Grimme et al., 2025; Grosswendt et al., 2014; Li et al., 2025; Manakov et al., 2022; Moore et al., 2015; Sheng et al., 2023).

## Contributions

L.L. designed and implemented the CLASHub platform, including web development, database integration, and analysis workflows, performed CLASH experiments, and wrote the manuscript. P.S., N.M.H., T.L., and Y.W. also performed CLASH experiments and contributed to data acquisition. A.L.G. assisted with interpretation of *C. elegans* CLASH. N.M.H. and C.M.T. provided feedback and minor revisions to the manuscript. M.X. supervised the project, provided overall guidance and co-drafted the manuscript. All authors contributed to data interpretation and approved the final version of the manuscript.

## Acknowledgements

We thank Dr. Bently P. Doonan for generously providing the human liver sample (IRB202002809). We acknowledge BioRender.com for assisting in the creation of the graphical abstract and user interface figures. We also used OpenAI ChatGPT-4.0 to assist with debugging and refining the HTML, JavaScript, CSS, and sbatch scripts for the CLASHub platform.

## Declaration of generative AI and AI-assisted technologies in the writing process

During the preparation of this work, the author(s) used OpenAI ChatGPT-4.0 in order to improve grammar and readability of the manuscript. After using this tool, the author(s) reviewed and edited the content as needed and take(s) full responsibility for the content of the publication.

---

**Figure S1. Overview of the number of samples and cell/tissue types represented in the CLASHub datasets across species.**

Bar plots summarizing the total number of samples (A) and total number of cell/tissue types (B) derived from published and newly generated CLASH data. Bar plots showing the number of unique cell and tissue types included in the CLASH (C), gene expression (D) and miRNA expression (E) data for human, mouse, *Drosophila*, and *C. elegans*, separated into control and ZSWIM8 knockout conditions.

**Figure S2. Example external resources linked from the CLASH results table.**

(A) miRBase page opened by clicking on the miRNA name (hsa-miR-7-5p). (B) Ensembl gene summary page opened by clicking on the gene ID (*OIP5-AS1*). (C) UCSC Genome Browser view showing the genomic locus corresponding to the hybrid site.

**Figure S3. CLASH analysis workflow for pre-processed FASTA inputs.**

(A) Interface for submitting CLASH analysis with cleaned FASTA files. Users (1) select the CLASH module, (2) choose “cleaned FASTA.gz” as input file type, (3) select a local FASTA file for upload, (4) select species, (5) specify output filename, (6) enter email address, and (7) submit the request. The backstage bioinformatic pipeline to process cleaned FASTA data is shown on the right.

**Figure S4. miRNA-seq analysis interface and processing overview.**

(A) Interface for miRNA-seq analysis. Users (1) select the module, (2) choose input file type, (3) select paired-end FASTQ files to upload, (4-5) enter adapter sequences, (6) specify output filename, (7) can optionally add additional samples, (8) select species, (9) provide email address, and (10) submit the request. The backstage bioinformatic pipeline to process paired-end FASTQ data is shown on the right.

(B) Output summary and a bar chart displaying statistics of the processed reads.

**Figure S5. miRNA-seq analysis workflows for single-end FASTQ and cleaned FASTA inputs.**

(A) Interface for processing single-end FASTQ files. Users (1) select the miRNA-seq module, (2) choose the “Single-End FASTQ” input, (3) choose a local file to upload, (4) enter the adapter sequence, (5) specify the output filename, (6) can optionally add additional samples, (7) select the species, (8) provide an email address, and (9) submit the request.

(B) Interface for processing cleaned FASTA files. Users (1) select the miRNA-seq module, (2) choose the “Cleaned FASTA” input, (3) choose a local file to upload, (4) specify the output filename, (5) can optionally add additional samples, (6) select the species, (7) provide an email address, and (8) submit.

**Figure S6. RNA-seq analysis module interface and processing summary.**

(A) Interface for RNA-seq gene expression quantification. Users (1) select the RNA-seq module, (2) choose paired-end FASTQ input, (3) upload Read 1 and Read 2 files, (4-5) enter adapter sequences, (6) specify output filename, (7) add more samples if needed, (8) select species, (9) provide email, and (10) submit the request. The backstage bioinformatic pipeline to process paired- end FASTQ data is shown on the right.

(B) Example processing output showing adapter sequences, read counts across each step, and a bar chart summarizing total, trimmed, and aligned reads.

**Figure S7. RNA-seq differential expression analysis module interface and workflow.**

Interface for differential gene expression analysis. Users (1) select the RNA-seq module, (2) choose the differential expression option, (3-10) upload at least two control group paired-end FASTQ files with the option to add additional samples (11), (12–20) upload two or more treatment group files, specify adapter sequences and output filenames, (21) select species, (22) provide email, and (23) submit the request. The backstage pipeline that performs adapter trimming, genome mapping, count generation, and differential expression analysis is shown on the right.

**Figure S8. Conservation and genome editing of the putative miR-335-3p trigger within *ATP6V1G1*.**

(A) UCSC Genome Browser view of the *Atp6v1g1* locus on mouse chromosome 4 (mm39), showing phyloP conservation scores across 35 vertebrates. The miR-335-3p binding site displays contiguous pairing at the seed and 3′ end regions.

(B) Schematic of the human *ATP6V1G1* transcript with the miR-335-3p TDMD trigger indicated (red arrow). Sanger sequencing results of CRISPR/Cas9-edited MDA-MB-231 clones are shown: M1 (108 bp deletion +1 bp insertion), M5 (118 bp deletion +6 bp insertion), and M8 (108 bp deletion).

(C) Schematic of the mouse *Atp6v1g1* transcript with the miR-335-3p TDMD trigger (red arrow). Sanger sequencing confirms four MEF clones (b5, b6, b11, b17) carrying identical 65 bp deletions.

