## Supplementary Figures for "CLASHub: an integrated database and analytical platform for microRNA-target interactions"

### Figure S1

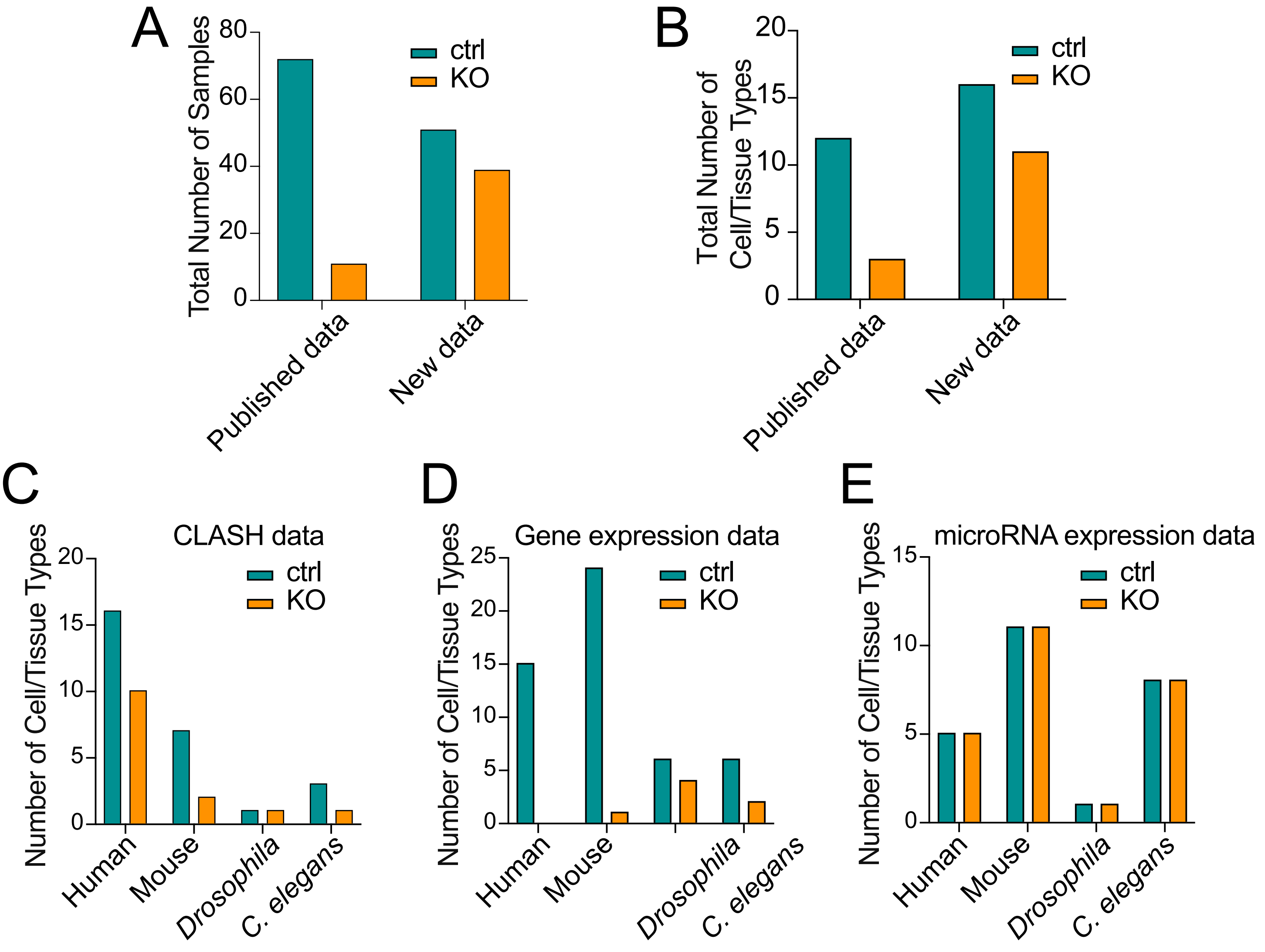
A
Total Number of Samples
80
60
40
20
0
ctrl
KO
Published data
New data
B
Total Number of Cell/Tissue Types
20
15
10
5
0
ctrl
KO
Published data
New data
C
CLASH data
Number of Cell/Tissue Types
20
15
10
5
0
ctrl
KO
Human
Mouse
Drosophila
C. elegans
D
Gene expression data
Number of Cell/Tissue Types
25
20
15
10
5
0
ctrl
KO
Human
Mouse
Drosophila
C. elegans
E
microRNA expression data
Number of Cell/Tissue Types
15
10
5
0
ctrl
KO
Human
Mouse
Drosophila
C. elegans

### Figure S2

A

Your query “**hsa-miR-7-5p**” returned **1** results

Download CSV File

| Section | #hits |
|---|---|
| Precursor miRNA | 0 |
| Mature miRNA | 1 |
| Dead | 0 |
| Previous ID | 0 |
| Dead entry previous ID | 0 |
| Previous mature ID | 0 |
| Gene symbols | 0 |
| **All matches** | **1** |

Show 10 entries

Filter:

| Name | Accession | Confidence |
|---|---|---|
| hsa-miR-7-5p | MIMAT0000252 | |

Showing 1 to 1 of 1 entries

Previous 1 Next

Select all | Reset | Fetch sequences

B

**Human** (GRCh38.p14)

**Gene: OIP-AS1 ENSG00000247556**

| | |
|---|---|
| **Description** | OIP5 antisense RNA 1 [Source:HGNC Symbol;Acc:HGNC:43563] |
| **Gene Synonyms** | CYRANO, LINC-OIP5 |
| **Location** | Chromosome 15: 41,283,958-41,322,392 forward strand. GRCh38:CM000677.2 |
| **About this gene** | This gene has 73 transcripts (splice variants). |
| **Gene Synonyms** | CYRANO, LINC-OIP5 |

C

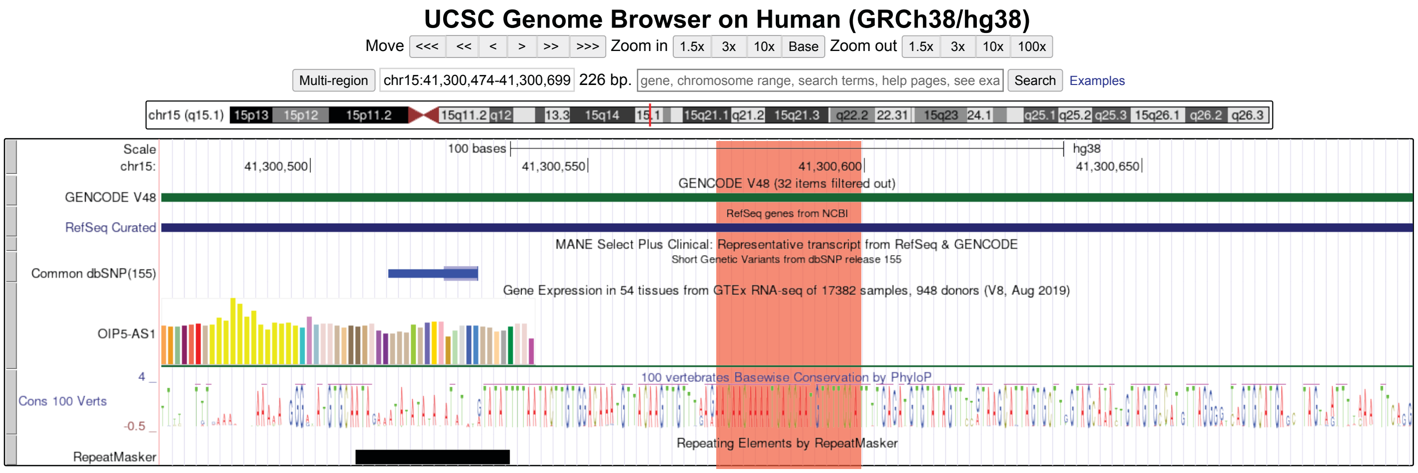

### Figure S3

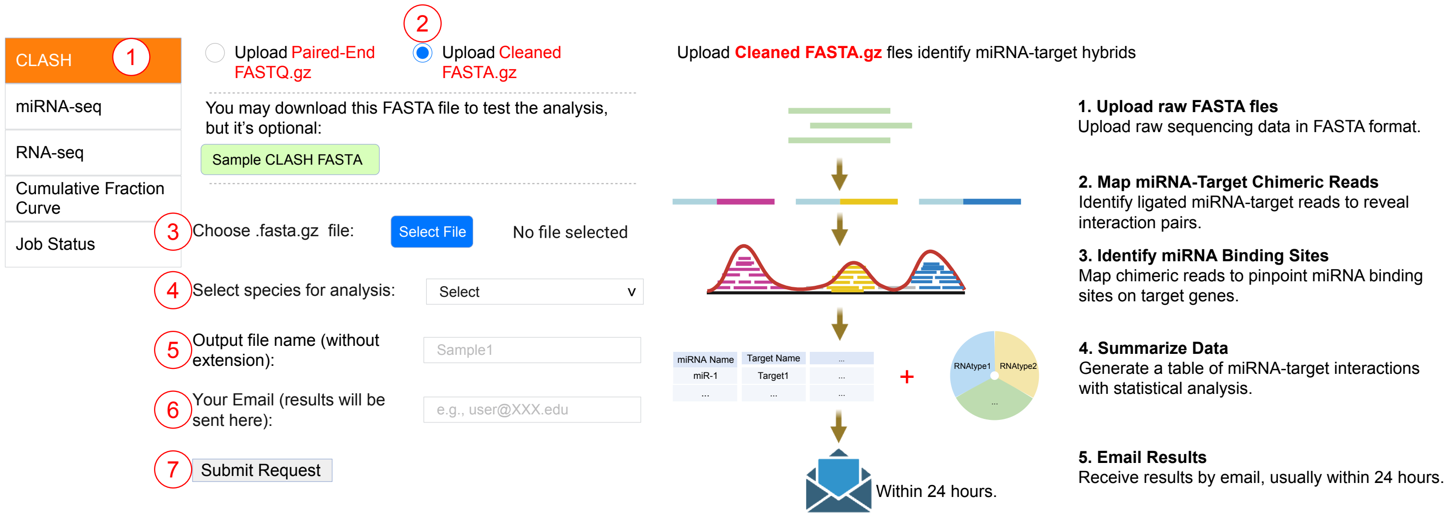
CLASH
1
miRNA-seq
RNA-seq
Cumulative Fraction Curve
Job Status
Upload Paired-End FASTQ.gz
2
Upload Cleaned FASTA.gz
You may download this FASTA file to test the analysis, but it's optional:
Sample CLASH FASTA
3
Choose .fasta.gz file:
Select File
No file selected
4
Select species for analysis:
Select
5
Output file name (without extension):
Sample1
6
Your Email (results will be sent here):
e.g.,
7
Submit Request
Upload Cleaned FASTA.gz fles identify miRNA-target hybrids
miRNA Name
Target Name
miR-1
Target1
RNAtype1
RNAtype2
Within 24 hours.
1. Upload raw FASTA fles
Upload raw sequencing data in FASTA format.
2. Map miRNA-Target Chimeric Reads
Identify ligated miRNA-target reads to reveal interaction pairs.
3. Identify miRNA Binding Sites
Map chimeric reads to pinpoint miRNA binding sites on target genes.
4. Summarize Data
Generate a table of miRNA-target interactions with statistical analysis.
5. Email Results
Receive results by email, usually within 24 hours.

### Figure S4

A

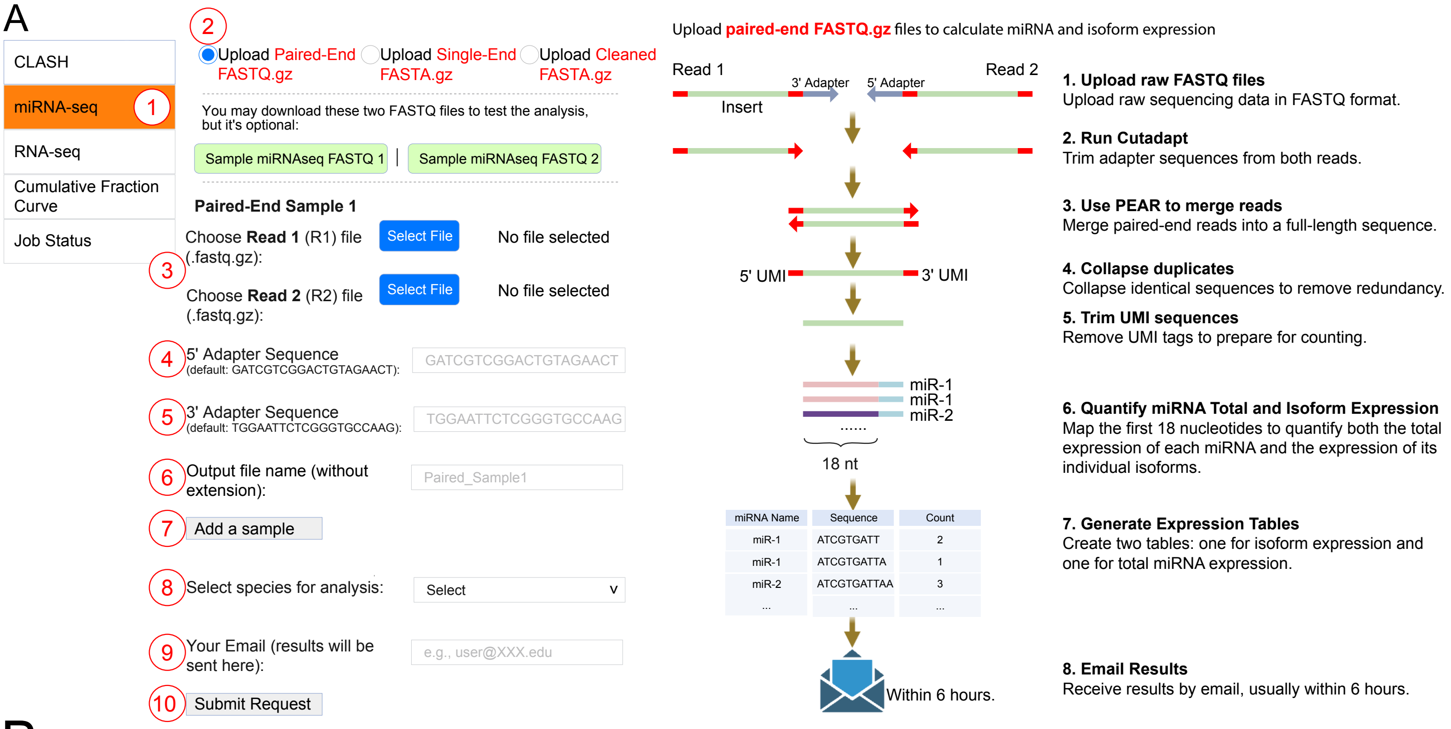


B

**Data Processing Summary**

| **Sample** | **Input File 1** | **Input File 2** |
|---|---|---|
| Sample 1 | Sample1_R1.fastq.gz | Sample2_R1.fastq.gz |
| 5' adapter | 3' Adapter | Total Reads |
| GATCGTCGGACTGTAGAACT | TGGAATTCTCGGGTGCCAAG | 25000 |
| Trimmed Reads | Collapsed Reads | Aligned Reads |
| 23479 | 20871 | 17400 |

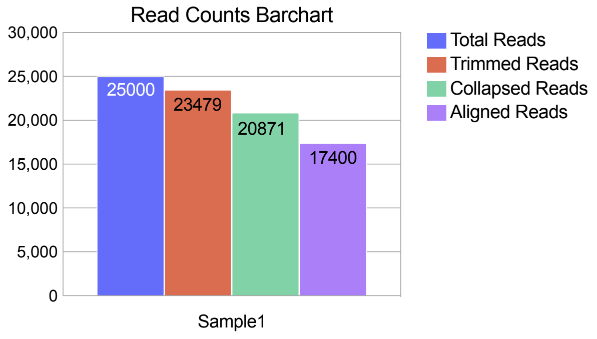

### Figure S5

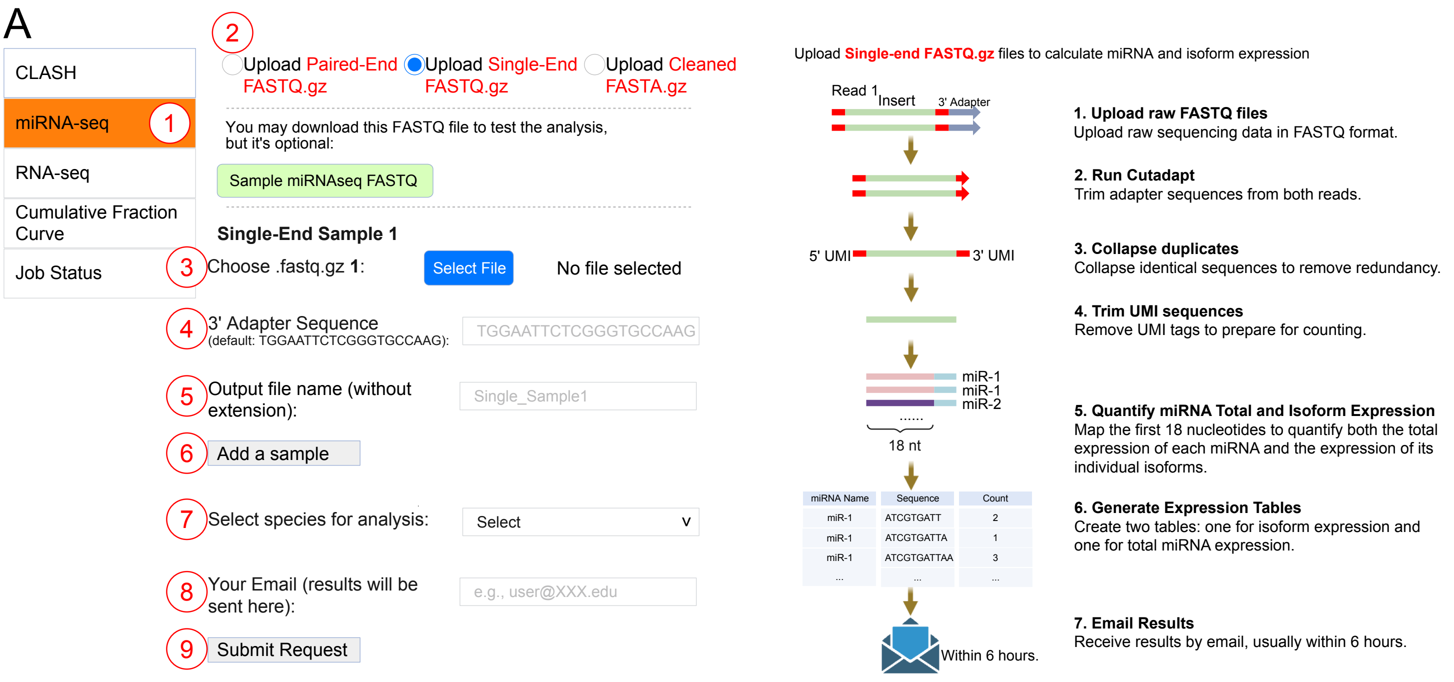
A
CLASH
miRNA-seq
RNA-seq
Cumulative Fraction Curve
Job Status
Upload Paired-End FASTQ.gz
Upload Single-End FASTQ.gz
Upload Cleaned FASTA.gz
You may download this FASTQ file to test the analysis, but it's optional:
Sample miRNAseq FASTQ
Single-End Sample 1
Choose .fastq.gz 1:
Select File
No file selected
3' Adapter Sequence
(default: TGGAATTCTCGGGTGCCAAG):
TGGAATTCTCGGGTGCCAAG
Output file name (without extension):
Single_Sample1
Add a sample
Select species for analysis:
Select
Your Email (results will be sent here):
e.g.,
Submit Request
Upload Single-end FASTQ.gz files to calculate miRNA and isoform expression
Read 1
Insert
3' Adapter
5' UMI
3' UMI
miR-1
miR-1
miR-2
18 nt
miRNA Name | Sequence | Count
miR-1 | ATCGTGATT | 2
miR-1 | ATCGTGATTA | 1
miR-1 | ATCGTGATTAA | 3
Within 6 hours.
1. Upload raw FASTQ files
Upload raw sequencing data in FASTQ format.
2. Run Cutadapt
Trim adapter sequences from both reads.
3. Collapse duplicates
Collapse identical sequences to remove redundancy.
4. Trim UMI sequences
Remove UMI tags to prepare for counting.
5. Quantify miRNA Total and Isoform Expression
Map the first 18 nucleotides to quantify both the total expression of each miRNA and the expression of its individual isoforms.
6. Generate Expression Tables
Create two tables: one for isoform expression and one for total miRNA expression.
7. Email Results
Receive results by email, usually within 6 hours.

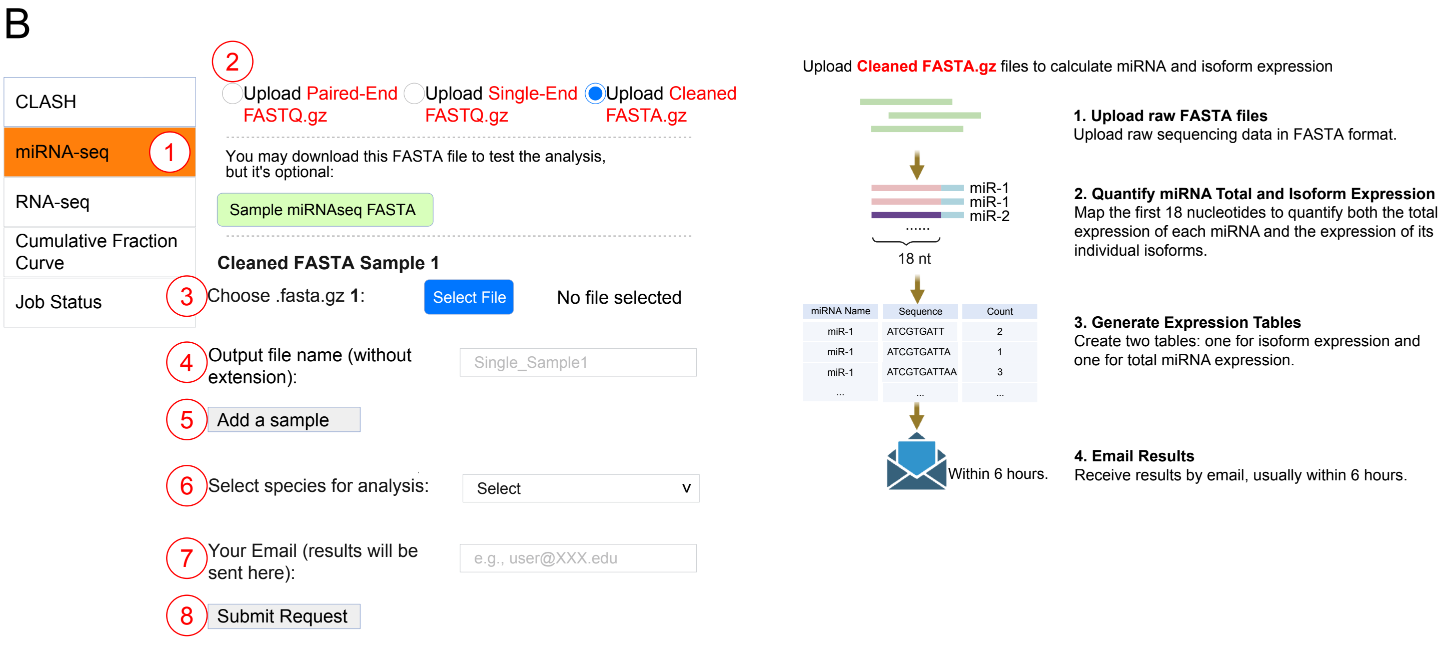
B
CLASH
miRNA-seq
RNA-seq
Cumulative Fraction Curve
Job Status
Upload Paired-End FASTQ.gz
Upload Single-End FASTQ.gz
Upload Cleaned FASTA.gz
You may download this FASTA file to test the analysis, but it's optional:
Sample miRNAseq FASTA
Cleaned FASTA Sample 1
Choose .fasta.gz 1:
Select File
No file selected
Output file name (without extension):
Single_Sample1
Add a sample
Select species for analysis:
Select
Your Email (results will be sent here):
e.g.,
Submit Request
Upload Cleaned FASTA.gz files to calculate miRNA and isoform expression
miR-1
miR-1
miR-2
18 nt
miRNA Name | Sequence | Count
miR-1 | ATCGTGATT | 2
miR-1 | ATCGTGATTA | 1
miR-1 | ATCGTGATTAA | 3
Within 6 hours.
1. Upload raw FASTA files
Upload raw sequencing data in FASTA format.
2. Quantify miRNA Total and Isoform Expression
Map the first 18 nucleotides to quantify both the total expression of each miRNA and the expression of its individual isoforms.
3. Generate Expression Tables
Create two tables: one for isoform expression and one for total miRNA expression.
4. Email Results
Receive results by email, usually within 6 hours.

### Figure S6

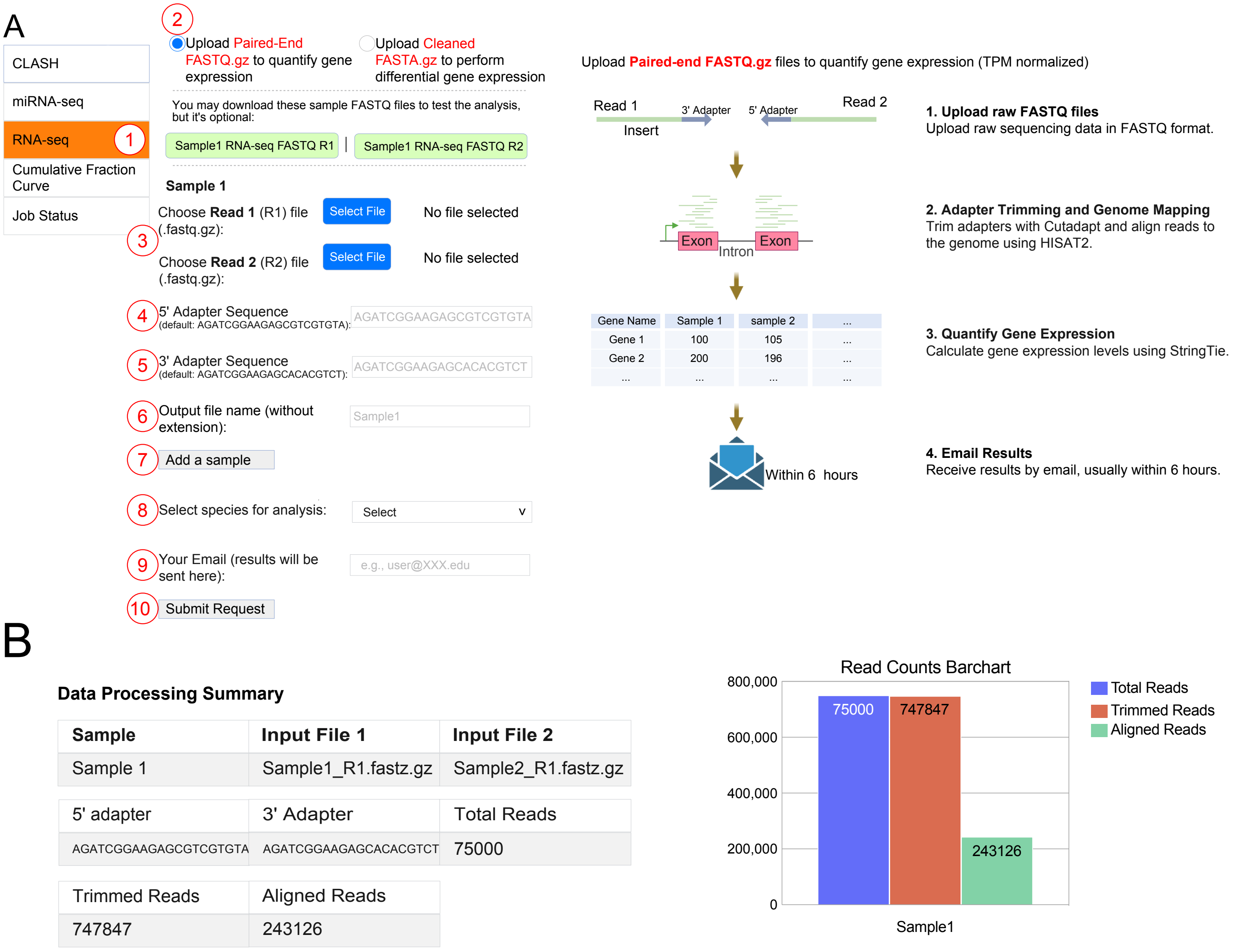
A
CLASH
miRNA-seq
RNA-seq
Cumulative Fraction Curve
Job Status
Upload Paired-End FASTQ.gz to quantify gene expression
Upload Cleaned FASTA.gz to perform differential gene expression
You may download these sample FASTQ files to test the analysis, but it's optional:
Sample1 RNA-seq FASTQ R1 | Sample1 RNA-seq FASTQ R2
Sample 1
Choose Read 1 (R1) file (.fastq.gz):
Select File
No file selected
Choose Read 2 (R2) file (.fastq.gz):
Select File
No file selected
5' Adapter Sequence (default: AGATCGGAAGAGCGTCGTGTA):
AGATCGGAAGAGCGTCGTGTA
3' Adapter Sequence (default: AGATCGGAAGAGCACACGTCT):
AGATCGGAAGAGCACACGTCT
Output file name (without extension):
Sample1
Add a sample
Select species for analysis:
Select
Your Email (results will be sent here):
e.g.,
Submit Request
Upload Paired-end FASTQ.gz files to quantify gene expression (TPM normalized)
Read 1
3' Adapter
5' Adapter
Read 2
Insert
Exon
Intron
Exon
Gene Name | Sample 1 | sample 2 | ...
Gene 1 | 100 | 105 | ...
Gene 2 | 200 | 196 | ...
Within 6 hours
1. Upload raw FASTQ files
Upload raw sequencing data in FASTQ format.
2. Adapter Trimming and Genome Mapping
Trim adapters with Cutadapt and align reads to the genome using HISAT2.
3. Quantify Gene Expression
Calculate gene expression levels using StringTie.
4. Email Results
Receive results by email, usually within 6 hours.
B
Data Processing Summary
Sample | Input File 1 | Input File 2
Sample 1 | Sample1_R1.fastz.gz | Sample2_R1.fastz.gz
5' adapter | 3' Adapter | Total Reads
AGATCGGAAGAGCGTCGTGTA | AGATCGGAAGAGCACACGTCT | 75000
Trimmed Reads | Aligned Reads
747847 | 243126
Read Counts Barchart
800,000
600,000
400,000
200,000
0
75000
747847
243126
Sample1
Total Reads
Trimmed Reads
Aligned Reads

### Figure S7

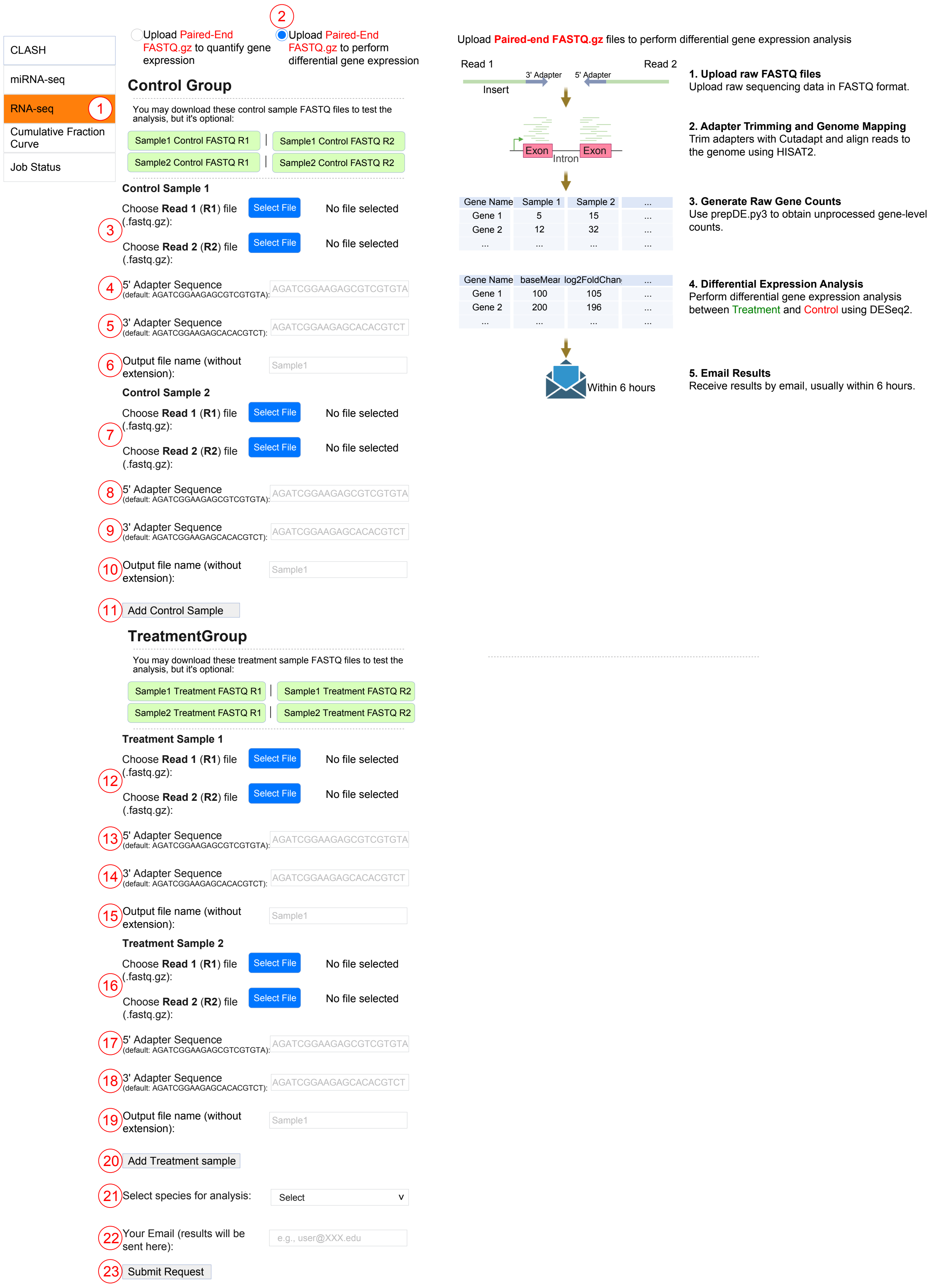

CLASH
miRNA-seq
RNA-seq
Cumulative Fraction Curve
Job Status
Upload Paired-End FASTQ.gz to quantify gene expression
Upload Paired-End FASTQ.gz to perform differential gene expression
Control Group
You may download these control sample FASTQ files to test the analysis, but it's optional:
Sample1 Control FASTQ R1 | Sample1 Control FASTQ R2
Sample2 Control FASTQ R1 | Sample2 Control FASTQ R2
Control Sample 1
Choose Read 1 (R1) file (.fastq.gz):
Select File
No file selected
Choose Read 2 (R2) file (.fastq.gz):
5' Adapter Sequence (default: AGATCGGAAGAGCGTCGTGTA):
AGATCGGAAGAGCGTCGTGTA
3' Adapter Sequence (default: AGATCGGAAGAGCACACGTCT):
AGATCGGAAGAGCACACGTCT
Output file name (without extension):
Sample1
Control Sample 2
Add Control Sample
TreatmentGroup
You may download these treatment sample FASTQ files to test the analysis, but it's optional:
Sample1 Treatment FASTQ R1 | Sample1 Treatment FASTQ R2
Sample2 Treatment FASTQ R1 | Sample2 Treatment FASTQ R2
Treatment Sample 1
Treatment Sample 2
Add Treatment sample
Select species for analysis:
Select
Your Email (results will be sent here):
e.g.,
Submit Request
Upload Paired-end FASTQ.gz files to perform differential gene expression analysis
Read 1
Read 2
3' Adapter
5' Adapter
Insert
Exon
Intron
Exon
Gene Name | Sample 1 | Sample 2
Gene 1 | 5 | 15
Gene 2 | 12 | 32
Gene Name | baseMean | log2FoldChange
Gene 1 | 100 | 105
Gene 2 | 200 | 196
Within 6 hours
1. Upload raw FASTQ files
Upload raw sequencing data in FASTQ format.
2. Adapter Trimming and Genome Mapping
Trim adapters with Cutadapt and align reads to the genome using HISAT2.
3. Generate Raw Gene Counts
Use prepDE.py3 to obtain unprocessed gene-level counts.
4. Differential Expression Analysis
Perform differential gene expression analysis between Treatment and Control using DESeq2.
5. Email Results
Receive results by email, usually within 6 hours.

### Figure S8

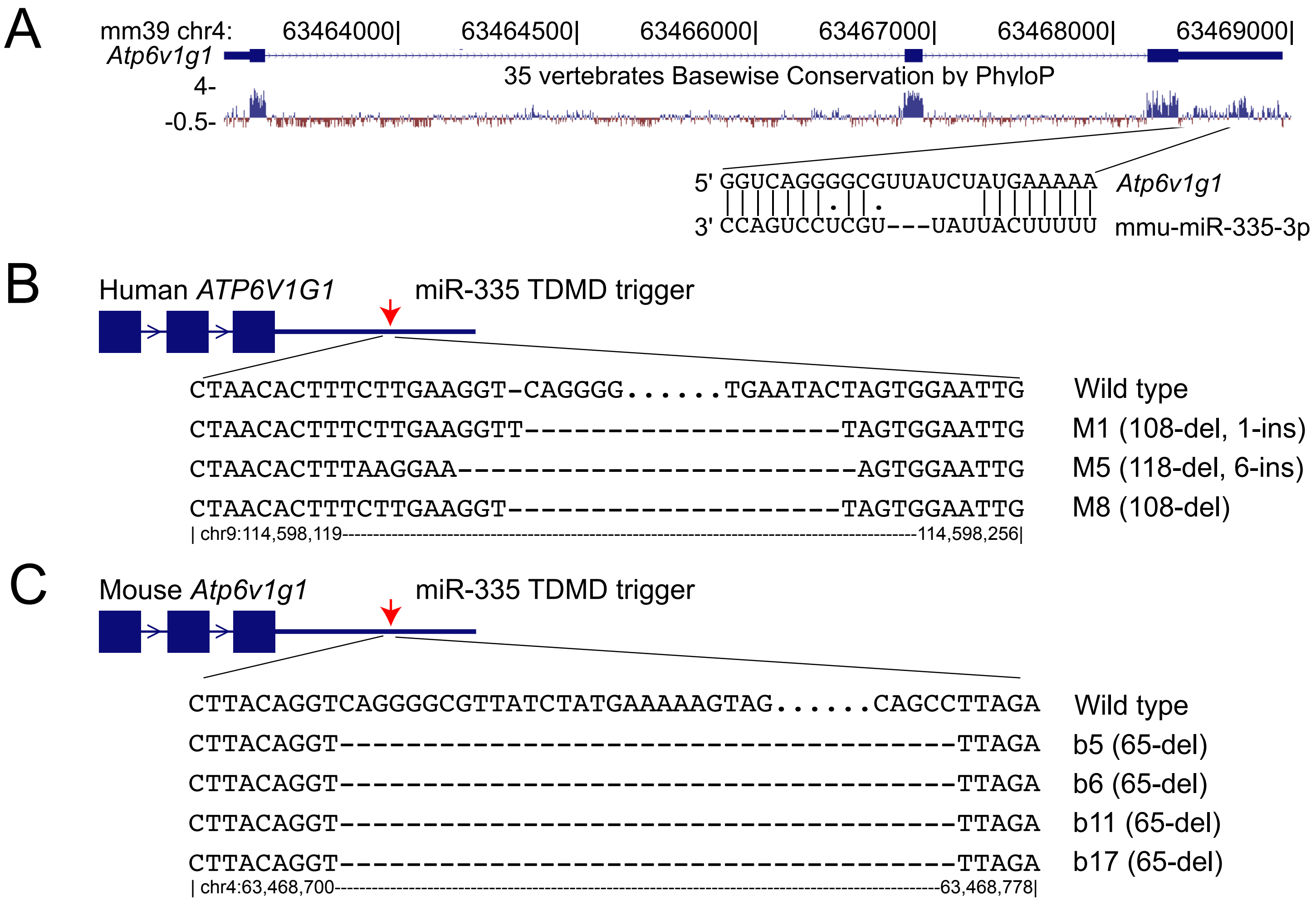
A
mm39 chr4: 63464000| 63464500| 63466000| 63467000| 63468000| 63469000|
Atp6v1g1
35 vertebrates Basewise Conservation by PhyloP
4-
-0.5-
5' GGUCAGGGGCGUUAUCUAUGAAAAA Atp6v1g1
3' CCAGUCCUCGU---UAUUACUUUUU mmu-miR-335-3p
B
Human ATP6V1G1 miR-335 TDMD trigger
CTAACACTTTCTTGAAGGT-CAGGGG......TGAATACTAGTGGAATTG Wild type
CTAACACTTTCTTGAAGGTT--------------------TAGTGGAATTG M1 (108-del, 1-ins)
CTAACACTTTAAGGAA-------------------------AGTGGAATTG M5 (118-del, 6-ins)
CTAACACTTTCTTGAAGGT---------------------TAGTGGAATTG M8 (108-del)
| chr9:114,598,119---114,598,256|
C
Mouse Atp6v1g1 miR-335 TDMD trigger
CTTACAGGTCAGGGGCGTTATCTATGAAAAAGTAG......CAGCCTTAGA Wild type
CTTACAGGT---------------------------------------TTAGA b5 (65-del)
CTTACAGGT---------------------------------------TTAGA b6 (65-del)
CTTACAGGT---------------------------------------TTAGA b11 (65-del)
CTTACAGGT---------------------------------------TTAGA b17 (65-del)
| chr4:63,468,700---63,468,778|
